# Pathogen-specific antimicrobial activity prediction with biological large language model-based methods

**DOI:** 10.64898/2026.05.23.727347

**Authors:** Berke Ucar, Emre Demirsoy, Ali Salehi, Darcy Sutherland, Anat Yanai, Lauren Coombe, Vanessa C. Thompson, René L. Warren, Caren C. Helbing, Inanc Birol

## Abstract

Driven by the rise of antimicrobial resistance, antimicrobial peptides (AMPs) have emerged as promising therapeutics capable of targeting multidrug-resistant pathogens. Because identifying AMPs and their specific targets requires costly and labor-intensive wet-lab experiments, *in silico* methods to prioritize candidates are highly valuable. However, current computational methods often lack pathogen specificity or fail to incorporate crucial targeted proteomic and genomic contexts. To bridge this gap, we developed triAMPh, a robust, zero-shot framework for pathogen-specific peptide bioactivity prediction. triAMPh integrates a heterogeneous graph attention network-based link predictor (HLP), Extreme Gradient Boosting, and a multilayer perceptron trained on features from biological large language models (bLLMs). Our novel HLP constructs a knowledge graph that maps peptides and pathogens as distinct nodes, connected by similarity and bioactivity edges. The model extracts information through semantic traversals, prioritizing neighboring nodes and their biological contexts. Benchmarking shows that triAMPh provides unbiased, peptide- and pathogen-centered zero-shot predictions, matching or outperforming state-of-the-art methods across all metrics except precision. Ultimately, triAMPh offers a powerful computational tool to accelerate wet-lab AMP discovery while demonstrating the capability of bLLMs to capture complex, pathogen-specific bioactivity patterns.

## INTRODUCTION

Antimicrobial peptides (AMPs) are increasingly recognized as promising therapeutic candidates to combat the escalating threat of AMR^1^. These evolutionarily conserved molecules are produced across all kingdoms of life as a part of their innate immune systems^2^ and exhibit wide-spectrum but target-specific activity against diverse biological targets, including bacteria, viruses, biofilms, parasites, and cancer cells^3^. An AMP can exert multiple mechanisms of action (MOAs) on relatively non-specific targets, hence, they are generally considered less prone to the development of AMR in target pathogens^2^. Additionally, unlike conventional antibiotics, AMPs primarily target the cell membrane and are thus often active against multidrug-resistant pathogens^2^.

The physicochemical properties of AMPs, such as charge density, amino acid composition, sequence length, three-dimensional confirmation, and hydrophobicity, are critical determinants of their bioactivity against different cells and molecular targets^4–8^. For example, variation in cell surface composition across different organisms shapes molecular interactions and determines the potency of membrane-targeting AMPs^9^. When large peptide libraries must be validated against different pathogens under limited resources to identify only a few promising candidates, even with improved throughput and reduced costs, purely wet-lab-based antimicrobial susceptibility testing (AST) methods remain relatively slow, expensive, and labour-intensive compared to hybrid approaches that combine experimental and computational screening^10^. Consequently, substantial advances in *in silico* approaches have recently been made to complement wet-lab screening methods for AMP discovery, supported by the increasing availability of AMP activity data^11^. Most of these methods tackle the problem on a binary (AMP vs. non-AMP)^12^, functional activity (e.g., antibacterial, antiviral) or target domain (e.g., anti-Gram-negative, anti-Gram-positive)^13^ level. However, as the activity of AMPs are dependent on the target pathogens^14^, there is a need for higher resolution in *in silico* predictions. But, as prediction resolution increases, challenges such as limited data availability and the disproportionate representation of certain pathogens^12^ persist, leaving generalizable pathogen-specific bioactivity prediction an open problem.

Most state-of-the-art tools aim to improve the target resolution by modelling bioactivity against individual species or strains, often by restricting the input data to a single pathogen per classifier or regressor^15^. Although less common, existing tools can predict the bioactivity of AMPs for multiple targets per model, using multitask learning approaches^16^ or ontology-based representations of pathogens to guide species-or strain-level bioactivity prediction^17^. However, these models are restricted to a limited set of pathogens and entirely overlook their genomic information. Only a limited number of studies have incorporated the genetic makeup of pathogens into their prediction approaches with utilizing broad phylogenetic metrics, such as digital DNA-DNA hybridization^18^ or simple compositional features^19^, which yield untargeted measures of overall strain similarity rather than capturing the precise molecular context required for targeted AMP prediction.

Biological large language models (bLLMs) have demonstrated success in learning deep representations of complex semantic relationships, evolutionary constraints, and functional implications of sequential characteristics^20^. Specifically, protein language models (pLMs) have demonstrated success in peptide bioactivity^21^, toxicity^22^, generation^23^, and targeted efficacy^24^ prediction tasks. Concurrently, Graph Neural Networks (GNNs) have proven effective in AMP discovery with the usage of the three-dimensional structures of the peptides to predict bioactivity^25^ or toxicity^22^. Building on these advances, we present a novel and zero-shot proof computational framework that integrates pLMs with nucleotide language models (nLMs) and a knowledge graph neural network to predict pathogen-level antimicrobial efficacy, to our knowledge, for the first time. We utilized the Evolutionary Scale Modelling 2 (ESM2)^26^ and Nucleotide Transformer version 2 (NTv2)^27^ embeddings with the ensemble of Extreme Gradiant Boosting (XGBoost), multilayer perceptron (MLP) and a Heterogeneous Graph Attention Network (HAN)^28^-based Link Predictor (HLP) (Figure 1). Our ensemble model, triAMPh, performs comparably or statistically significantly better than state-of-the-art methods on two curated datasets (both public and in-house), the compilation of which constitutes a further contribution of this work.

**Figure 1.**
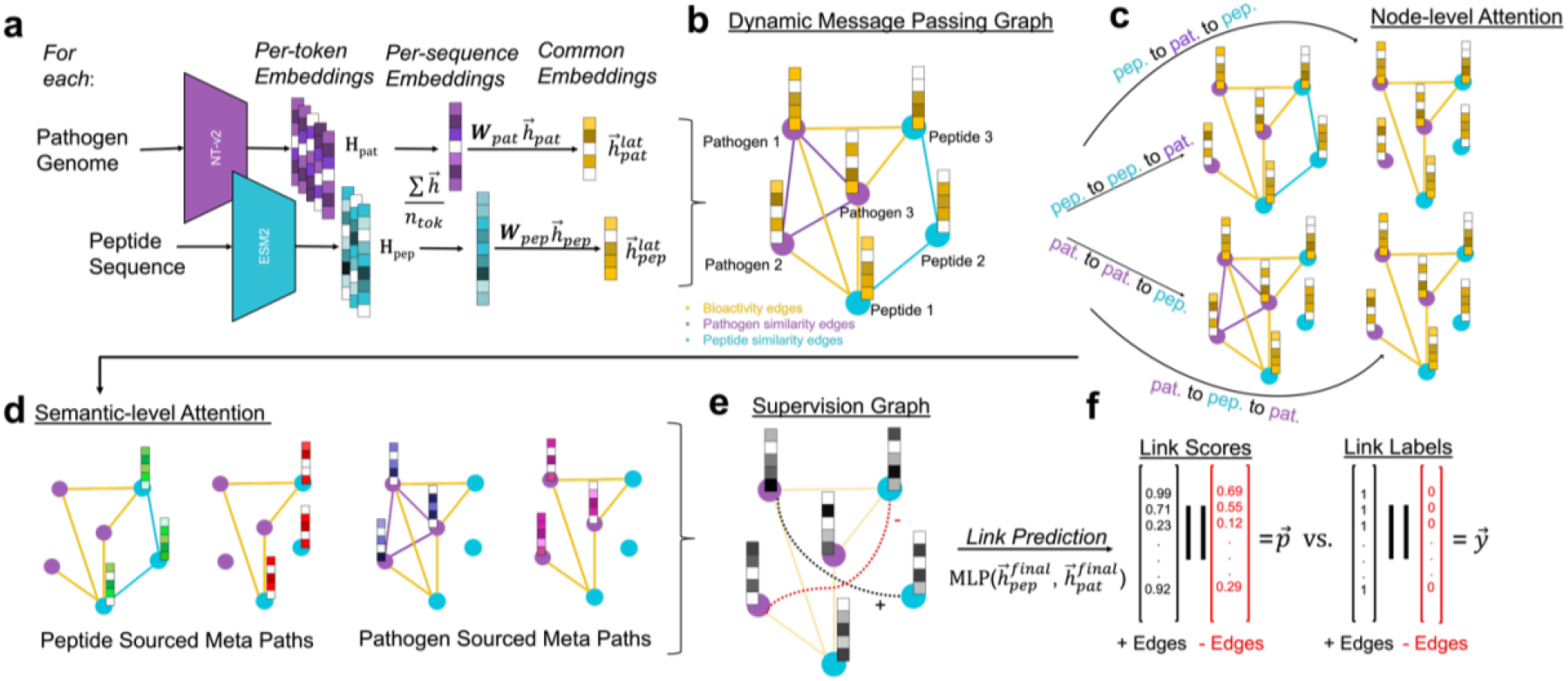
Heterogeneous Attention Network-based Link Predictor model architecture. Blue nodes and links represent peptide related nodes and edges. Purple nodes and links denote pathogen related nodes and links. **a**. bLLM embeddings are mean-pooled and projected into a shared representation space. **b**. A message-passing graph is then constructed dynamically, using pairwise distances between the embeddings and incorporating a subset of known peptide–pathogen bioactivity data. **c**. The message passing graph is deconstructed into distinct metapath–reachable subgraphs, each encoding a different type of relational context between neighbouring nodes. Node-level attention is applied within each subgraph to learn localized interactions. **d**. To integrate these diverse relational signals, semantic attention is used to aggregate the updated node embeddings across the different meta paths. Each vector colour means different semantics. **e**. A multilayer preceptron predicts interaction scores for previously unseen peptide– pathogen pairs based on their learned representations. These scores, alongside the predictions from other machine learning models, are then fed into a meta learner to form the triAMPh ensemble.

## RESULTS

### Assessing the triAMPh Ensemble

To justify the constituent models within the triAMPh ensemble, we evaluated their performance across two datasets: an in-house set and a mined test set. This mined test set served as the standard hold-out data from the total mined dataset, consisting of samples excluded during the training and validation phases. We compared individual models which establish triAMPh, a baseline Support Vector Machine model, and diverse ensemble of these four individual models. A Blending ensemble approach was employed, using a Logistic Regression meta learner. Figure 2 summarizes the performance and statistical comparison of four individual models and their 11 different combinations.

**Figure 2.**
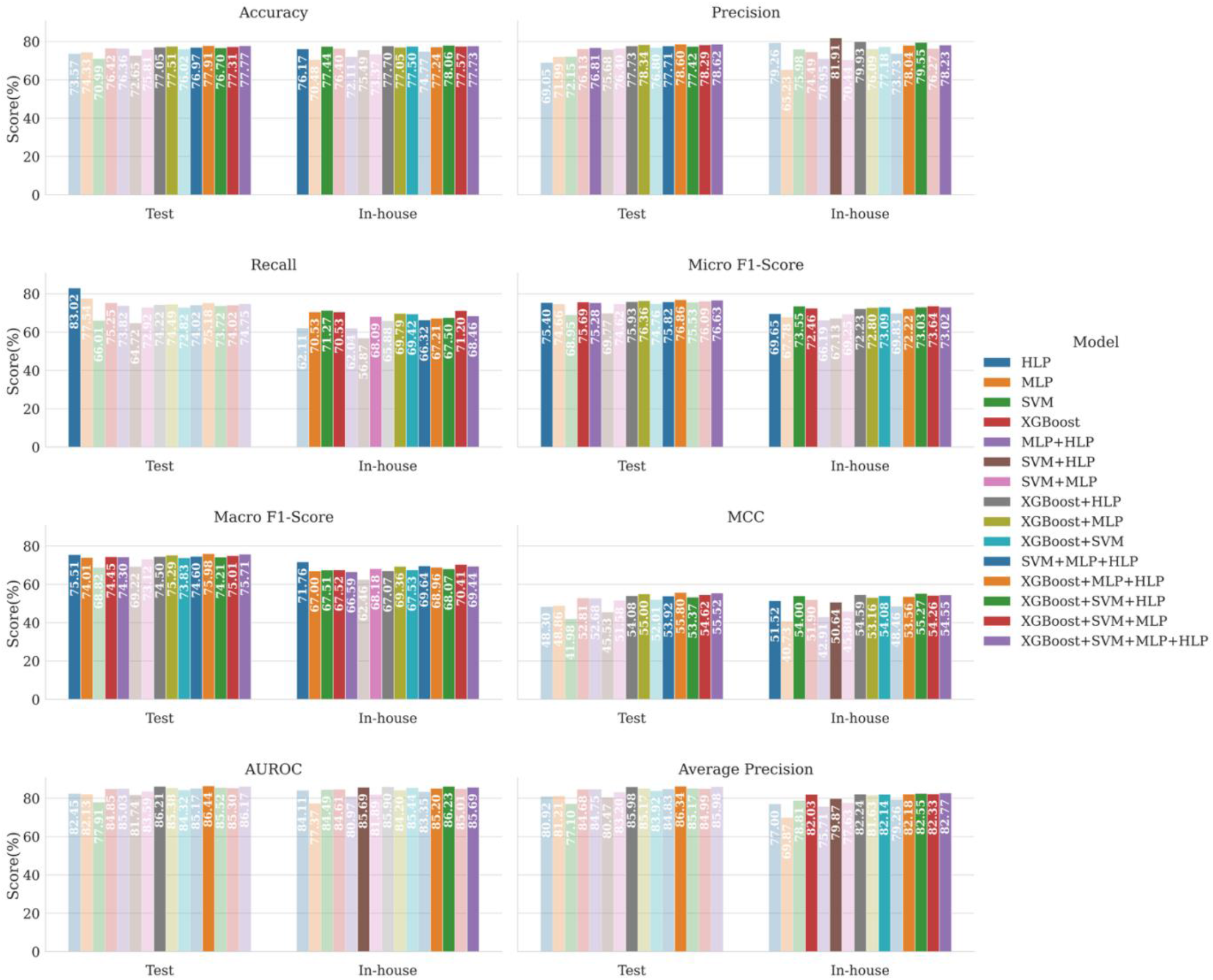
Performance on the Inductive individual and ensemble models on two of the test sets. The mined test dataset was comprised of 6,171 bioactivity pairs, while in-house dataset contained 3,076 bioactivity pairs. Transparency denotes statistical significance (p-value < 0.01) in the performance gap compared to the best model on the same dataset based on 100,000 bootstrapping iterations followed by Holm-Bonferroni correction.

Evaluating across two datasets, the triAMPh ensemble demonstrated the highest success rate among all evaluated methods, either significantly outperforming competitors or reaching statistical equivalence in 15 of 16 comparisons (eight metrics each for two datasets). Notably, for the sole metric where the ensemble did not match the top performer (recall on the mined test set), its constituent HLP module achieved parity on its own. Furthermore, the HLP module emerged as the most robust standalone system, attaining statistical equivalence with the best systems in seven distinct comparisons.

Because the primary use case of triAMPh is the targeted bioactivity inference of unseen peptides, we employed Inductive learning to train the individual constituent models and the ensemble, partitioning the data by unique peptides (Methods). To empirically validate this choice, we evaluated the performance of individual model, including the SVM baseline, trained via both Transductive and Inductive learning on the mined test sets (Supplementary Figure 2) and the in-house set. On the mined test set (Supplementary Figure 3), the top-performing models (including statistical equivalences) were consistent across both learning schemes, with the exception of the micro F1-score. This indicates that universal performance of different architecture on the mined dataset was largely consistent and independent from the learning strategy. Conversely, on the in-house test set (Supplementary Figure 4), models trained using inductive learning consistently achieved the highest performance or statistical equivalence to the top performer. To investigate this further, we directly compared the performance of identical model architectures trained under the two different schemes. This analysis revealed that only Inductive variants demonstrated statistical superiority over their Transductive counterparts. Ultimately, because the in-house dataset consists predominantly of peptides previously unseen by the models, these findings verify that inductive learning is more effective for predicting the bioactivity of novel peptides.

In addition to triAMPh’s superior success rate, we observed that the performance of the individual models varied across datasets. Ultimately, these fluctuations underscore the necessity of an ensemble strategy, combining XGBoost, HLP, and MLP, to ensure robust, pathogen-specific bioactivity prediction. A further key takeaway is that, under the employed learning strategy, HLP emerged as the strongest standalone model, yielding results that most closely approximate the peak performance achieved by the full ensemble.

### Per-Species Performance of triAMPh

To assess potential biases in the predictions made by triAMPh, we evaluated its pathogen-specific performance on both the mined test and in-house datasets. Figure 3 summarizes the F1-score, relative proportion, ratio of positive to total samples, and positive prediction rate (PPR; defined as the ratio of positive predictions), stratified by the pathogens associated with the bioactivity pairs across the different datasets.

**Figure 3.**
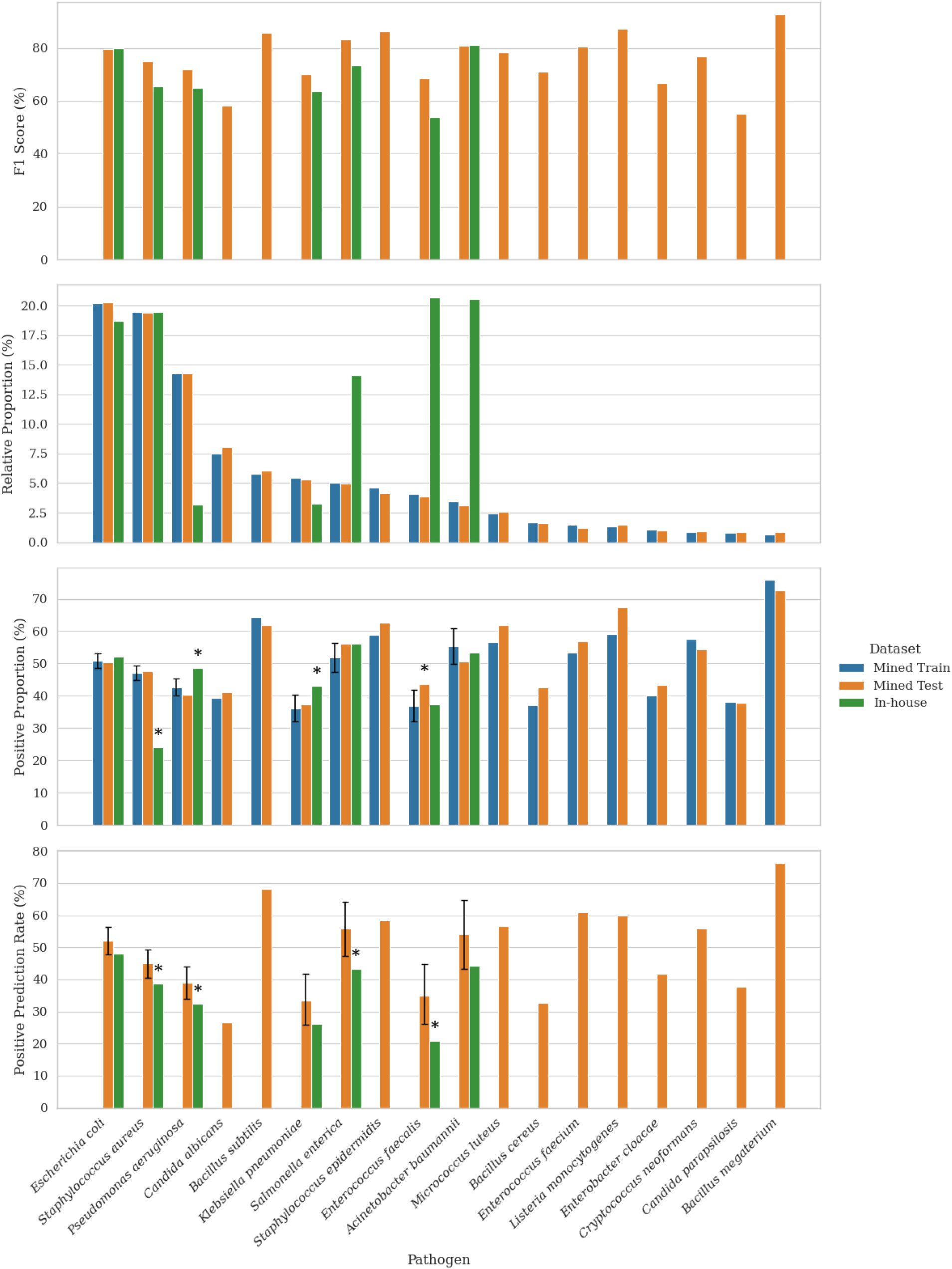
Performance metrics and dataset distribution of triAMPh across training and test sets. The columns are ordered based on the relative proportion of pathogen associated pairs within the training set. Error bars indicate 95% Wilson score intervals corrected with Bonferroni’s method, which were calculated only for pathogens included in both test sets. Asterisks denote statistically significant differences on columns compared to the column given with an error bar in the same pathogen.

The training set prevalence of peptide-pathogen pairs associated with a specific pathogen did not affect overall performance, as measured by the F1-score. Additionally, changes in prevalence relative to the training set had no apparent impact on prediction quality. For example, the performance observed for *Staphylococcus aureus*- and *Pseudomonas aeruginosa*-associated pairs was very similar across both sets, even though pairs associated with *P. aeruginosa* had a substantially lower prevalence in the in-house set than in the training set.

Additionally, we investigated the potential effect of data composition within pathogen stratified subsets on prediction bias. Here, we assumed that a composition biased model would predict the same rate of positives even when the proportion positive samples in the test set is significantly different than its training set. Among the four stratified subsets which had one of the test sets exhibiting significant composition deviation from the training set, only *Klebsiella pneumoniae* subsets’ PPRs on the test and in-house sets were not significantly different. *S. enterica*-associated PPRs on tested sets were significantly different, even though the sets compositions were statistically equivalent. We concluded that dataset composition did not affect the prediction capabilities of triAMPh. Even when positive sample distributions differed statistically across datasets, or when the PPR differed significantly despite equivalent positive proportions, the model did not default to predicting a consistent proportion of positives.

### triAMPh Provides Robust Pathogen-centric Zero-Shot Prediction Performance

Even though we anticipate the main use case of triAMPh will be predicting the bioactivity of unseen peptides against seen pathogens, we investigated whether evolutionary information encoded in nLM embeddings is indeed utilized by triAMPh, rather than being used as proxy labels for specific pathogens. We tested triAMPh on 1,861 bioactivity pairs associated with six pathogens that were not included in the training. These pathogens were the most prevalent six remaining within the mined databases. The pathogen stratified performance of triAMPh is represented in Table 1.

**Table 1.** Pathogen-centric zero-shot performance of triAMPh. All statistics are reported as percentages.

| Pathogen | Count | Accuracy | Precision | Recall | F1-Score | MCC | AUROC | Average Precision |
| --- | --- | --- | --- | --- | --- | --- | --- | --- |
| Overall | 1,861 | 66.79 | 81.79 | 63.95 | 71.78 | 34.37 | 73.90 | 80.92 |
| *Candida tropicalis* | 344 | 61.92 | 92.20 | 52.00 | 66.50 | 36.51 | 81.75 | 91.22 |
| *Streptococcus mutans* | 327 | 76.45 | 89.42 | 74.78 | 81.45 | 51.43 | 86.45 | 92.92 |
| *Streptococcus pyogenes* | 319 | 73.04 | 92.22 | 69.75 | 79.43 | 46.05 | 85.07 | 93.56 |
| *Saccharomyces cerevisiae* | 298 | 73.49 | 84.57 | 74.00 | 78.93 | 44.32 | 79.48 | 86.07 |
| *Pseudomonas syringae* | 289 | 51.90 | 96.75 | 46.85 | 63.13 | 23.37 | 67.53 | 93.30 |
| *Proteus mirabilis* | 284 | 62.68 | 35.29 | 88.52 | 50.47 | 36.36 | 80.84 | 52.19 |

Overall, we observed 69.98% macro F1-score on these predictions, comparable with both test and in-house dataset performance of triAMPh. Consistent with the previously observed per-species performance, triAMPh predictions did not show any trend associated with the size of the pathogen-related bioactivity pairs. Except for *Proteus mirabilis* predictions, the triAMPh predictions had higher precision and lower recall, suggesting more false negatives. This aligns with the results from the in-house test set, where the predicted peptide-pathogen pairs primarily involved unseen peptides.

### Comparing triAMPh against state-of-the-art methods

Two state-of-the-art methods, namely AMPSpeciesSpecific and Vishnepolsky et al.’s model, were tested on subsets of the inductive test set and the in-house set. Neither model offered the option for retraining. Hence, to make the comparisons as fair as possible, we took subsets of the test datasets based on each of the tools’ original training data constraints, pathogen set, and considerations for classifying peptide activity (Supplementary Methods). These datasets were used to benchmark the state-of-the-art methods against triAMPh. Tables 2 and 3 summarize the results of testing performed on the mined and in-house test sets, respectively.

**Table 2.** Benchmarking competitor models against the top-performing methods from the present study on respective subsets of the Mined Dataset. All results are presented as percentages, with the leading metric for each dataset highlighted in bold. For each dataset, the italicized values are significantly worse than the bolded ones. Vishnepolsky et al. provided a different prediction score than logits, hence AUROC and Average Precision cannot be calculated.

| Model | Dataset Details | Accuracy | Precision | Recall | Micro F1-Score | Macro F1-Score | MCC | AUROC | Average Precision |
| --- | --- | --- | --- | --- | --- | --- | --- | --- | --- |
| AMPSpecies Specific | 3,910 pairs, 5 species | *61.10* | *56.56* | **87.75** | *68.79* | *70.98* | *27.30* | *71.86* | *71.32* |
| triAMPh |  | **78.67** | **78.29** | *77.96* | **78.12** | **79.62** | **57.31** | **87.13** | **87.10** |
| Vishnepolsky et al. | 1,065 pairs, 8 strains | **83.47** | **82.68** | **84.25** | **83.46** | **84.32** | **66.96** | - | - |
| triAMPh |  | 81.41 | 79.85 | 83.49 | 81.63 | 80.46 | 62.89 | 89.58 | 89.92 |

**Table 3.** Benchmarking competitor models against the top-performing methods from the present study on respective subsets of the In-house Dataset. All results are presented as percentages, with the leading metric for each dataset highlighted in bold. For each dataset, the italicized values are significantly worse than the bolded ones. Vishnepolsky et al. provided a different prediction score than logits, hence AUROC and Average Precision cannot be calculated.

| Model | Dataset Details | Accuracy | Precision | Recall | Micro F1-Score | Macro F1-Score | MCC | AUROC | Average Precision |
| --- | --- | --- | --- | --- | --- | --- | --- | --- | --- |
| AMPSpecies Specific | 1,272 pairs, 3 species | *52.99* | *44.88* | **94.51** | *60.86* | 63.40 | *26.69* | *80.92* | 74.35 |
| triAMPh |  | **78.54** | **70.32** | *77.03* | **73.52** | **70.15** | **55.70** | **85.57** | **80.24** |
| Vishnepolsky et al. | 881 pairs, 4 strains | *72.87* | *65.35* | *52.37* | *58.14* | *53.91* | *38.95* | - | - |
| triAMPh |  | **83.09** | **80.00** | **70.66** | **75.04** | **76.64** | **62.60** | **88.53** | **84.57** |

Compared to AMPSpeciesSpecific, our ensemble model demonstrated significantly better performance, with improvements ranging from 9% to 30% across all metrics, with the exception of recall. While Vishnepolsky et al.’s model outperformed our ensemble, the differences in performance were not statistically significant. We hypothesize that the model’s slightly higher, yet statistically insignificant, performance was due to data leakage, as 412 of the 1,065 tested pairs were already present in its training set.

When using the in-house dataset, triAMPh outperformed both other methods across all metrics by approximately 4% to 24%, except for Recall compared to AMPSpeciesSpecific. Most of these improvements were statistically significant, except for the macro F1-score and average precision relative to AMPSpeciesSpecific. Overall, these results underscore the robustness of triAMPh, particularly when applied to data distributions distinct from those found in available public databases.

## DISCUSSION

Here, we present triAMPh, a framework that integrates XGBoost, MLP, and a novel HLP, built upon pretrained bLLM embeddings for pathogen-specific bioactivity inference. The HLP utilizes a two-stage attention mechanism that first captures neighborhood-based associations within a single metapath, and subsequently aggregates the extracted semantic-specific information based on metapath-level relationships. We trained triAMPh using a peptide-centric, zero-shot data splitting scheme, partitioning the dataset by unique peptides rather than peptide-pathogen pairs. To justify triAMPh’s computational complexity, we compared it against its individual constituent models, a baseline SVM, and ensemble combinations of these four models. triAMPh achieved the highest performance—or statistical equivalence to the top performer— in 15 out of 16 comparisons across eight metrics and two datasets. Overall, triAMPh achieved the highest number of top performances (including statistical equivalences) across these comparisons, with the next-best ensembles securing 13 out of 16.

Given its strong overall performance, we investigated the per-species accuracy of triAMPh to determine whether it exhibits any composition-related biases. triAMPh demonstrated no apparent correlation between its predictive performance and the number of pathogen-specific pairs included in the training set, indicating that high performance on more prevalent species is not a primary driver of its success. Additionally, we examined how the ratio of positive samples within the species-specific training data affected the model’s positive PPR. We found that triAMPh’s PPR was independent of the positive class distribution in the training dataset. This suggests that rather than defaulting to dataset label frequencies, triAMPh successfully captures the underlying mechanistic properties governing antimicrobial interaction patterns.

Beyond the peptide-centric, zero-shot evaluation, we tested triAMPh on publicly mined bioactivity pairs associated with six unseen pathogens. The framework achieved results comparable to those observed on the in-house dataset, demonstrating that even without explicit pathogen-centric optimization, triAMPh effectively extracts relevant signals from the nLM embeddings for pathogen-specific bioactivity prediction. Together, these robust peptide- and pathogen-centric zero-shot results indicate that bLLMs can capture underlying antimicrobial interaction patterns without the need for fine-tuning.

Finally, we compared triAMPh against two state-of-the-art pathogen-specific models on subsets of the mined and in-house test sets. In most cases, triAMPh demonstrated statistically significant improvements over these baselines. The only exception was its performance on the mined test set compared to the model by Vishnepolsky et al.; however, this anomaly is likely attributable to a severe data leak we detected between their training data and the test set. Of the two baselines, AMPSpeciesSpecific omits pathogen genetic information entirely, while the other relies on non-targeted phylogenetic distances. In contrast, by leveraging an nLM, triAMPh achieved significantly better or equivalent performance, demonstrating that nLMs can provide broad yet highly targeted insights into antimicrobial susceptibility.

A broader observation was that all evaluated models, including competitor baselines, performed substantially worse on the in-house test set compared to the mined test set. This drop occurred even though the optimization of the inductive models was consistent with zero-shot learning principles. Because our in-house dataset was generated through a novel AMP discovery pipeline, it comprises peptide-pathogen pairs that are highly distinct from those found in public resources, which inherently increases the difficulty of the prediction task. Despite this challenge, triAMPh significantly outperformed the other methods, demonstrating its robust generalization capabilities in zero-shot prediction scenarios.

Conventional functional inference methods rely on representations of amino acid sequences such as one-hot encoding^11^, compositional summaries^18^, and overall physicochemical properties^29^. However, these encodings do not represent biological effects of amino acids or disregard context in sequences. Furthermore, they necessitate cumbersome feature engineering and extensive prior knowledge of a peptide’s desired properties. These limitations similarly affect most applications involving targeted peptide bioactivity prediction, which also require investigation of genomic content and context. Most existing methods addressing the targeted bioactivity problem suffer from several limitations, such as restricting the prediction scope by excluding genomic content^16^, disregarding genomic context^30^, or utilizing overly broad comparative genomic measures^18^. triAMPh uniquely overcomes these challenges by leveraging bLLMs. We have provided empirical evidence that the joint application of pLMs and nnLMs is a superior approach compared to existing methodologies in the literature. Furthermore, these bLLM-based representations completely eliminate the need for manual feature engineering while capturing evolutionary and functional insights that align with antimicrobial activity patterns.

We acknowledge that this study is subject to several limitations. Because our approach relies on AST data, triAMPh only predicts peptide bioactivity with direct antimicrobial mechanisms, potentially overlooking indirect modes of action. Additionally, chromosomal DNA and plasmids were treated as linear sequences (which might not be the case for some organisms^31^) and merged during embedding generation. This strategy may introduce artifacts in cases where embeddings are generated across the junction of the chromosomal DNA and the plasmid within a single 1,000-token window. Our study also inherits the limitations of the used bLLMs. ESM2 was trained on the UniRef protein sequence database^32^, which compiles proteomes from across the tree of life but disproportionately overrepresents well-studied organisms. Similarly, the variant used for NTv2 was trained on a multi-species dataset that was heavily skewed toward vertebrate nucleotides.

Specifically, HLP utilizes an information diffusion scheme that relies on established bioactivity edges between peptides and pathogens. Consequently, a subset of known bioactivity pairs is required to construct the graph structure. This necessitates the maintenance of an auxiliary “knowledgebase” during training and inference, distinct from the supervision data. In the current study, we observed that HLP favoured positive predictions more on the mined test set for Transductive models, where the average number of connections per peptide in the message-passing graph was higher than that of the training set. Additionally, the Transductive models exhibited a higher false-negative rate on the in-house dataset, likely because the tested peptides lacked bioactivity connections within the message-passing graph (Supplementary Figures 3 and 4). Therefore, when applying the model to independent test sets, the underlying knowledgebase must be carefully tuned to ensure relevance.

Leveraging the rapidly growing volume of biological data can enhance both scope and granularity of this study. First, the dataset could be expanded to include a broader diversity of pathogens, such as viral species. As data density increases, the HLP framework can be evolved into a more comprehensive heterogeneous graph, incorporating strain-level nodes to enable higher-resolution predictions. Ideally, this would facilitate the training of strain-specific MIC predictors using bLLM embeddings. Also, the heterogeneous knowledge graph could be utilized to identify synergistic or antagonistic peptide interactions. Finally, the pathogen-specific bioactivity predictor holds significant potential as a guidance system for generative protein models, enabling the *de novo* design of AMPs with targeted bioactivity.

triAMPh is a state-of-the-art, zero-shot, pathogen-specific bioactivity prediction framework. To our knowledge, it is the first model in the AMP discovery domain to use a combination of pLMs and nLMs, as well as knowledge network representations of peptides and pathogens. It offers a practical and state-of-the-art solution to streamline the discovery process with assisting wet-lab bioactivity assessment. Broadly, this research offers empirical evidence that pre-trained bLLMs can be successfully deployed across various domains without requiring task-specific fine-tuning. We hope this work encourages the synergistic application of bLLMs in future functional proteomics research.

## METHODS

### Public Dataset Mining

We mined three publicly available AMP databases that provide experimental MIC at, at least, the species-level resolution: DBAASP v3^33^, dbAMP v3^34^, and DRAMP 4.0^35^ (accessed in July, released in June, and November 2024, respectively). Only measurements reported as MICs (i.e., filtering out MIC50, MIC90, etc.) were included to enable a fair comparison. Peptides that contain non-proteinogenic amino acids, are non-monomeric, or longer than 1,022 amino acids, were removed from the dataset due to ESM2’s constraints. Additionally, reported PTMs were disregarded, due to constraints of ESM2. Furthermore, experimental conditions for MIC measurements were excluded due to the lack of standardization across studies, which would have otherwise severely limited the sample size. Since databases contained different ways of reporting MICs, i.e. as an exact value or an interval, we used a set of criteria to standardize the representation, as summarized in Supplementary Table 1. These criteria were mainly based on the geometric average rather than the arithmetic, as MICs are conventionally measured in a twofold dilution series^36^.

We then standardized all measurements to micromolar (µM), as molecule density directly influences peptide–cell interactions^37^. Using molar concentration eliminates biases introduced by peptide molecular weight or length, since it reflects the actual number of molecules present in a given volume. Following this conversion, the dataset was divided into positive and negative subsets using a threshold of 32 µM. For each species, if any MIC value reported for a peptide against any strain of that species was less than or equal to the threshold, the peptide–species pair was assigned to the positive subset and removed from the negative subset. Peptides-pathogen pairs that were reported with an MIC value containing “>” in the original occurrence, which then turned out to be smaller than or equal to our threshold value, were not included in the positive subset. In the literature, peptides exhibiting MIC values exceeding 128 µg/mL are typically classified as inactive^38,39^. Given the mean molecular weight of the unique peptides identified (∼2268.31 g/mol), this concentration corresponds to approximately 56.43 µM. To be conservative with the bioactivity classification, the largest power of two below this value, 32µM, was defined as the threshold for distinguishing between positive and negative sets.

Duplicate peptide–species entries were then removed within each subset. Finally, species with more than 350 unique peptides reported as either active or inactive were retained in the final dataset. Here, we used the abbreviated genus names to filter for pathogens. A threshold of 350 peptides tested per pathogen was selected to optimize the trade-off between pathogen diversity and prevalence. The final dataset comprises 30,069 positive and 16,130 negative bioactivity peptide-pathogen pairs, collectively containing 11,741 unique peptides and 18 unique species (Supplementary Figure 1). The curated database of peptide-pathogen pairs is referred to as mined dataset. From the remaining datapoints, the next most prevalent six pathogens were selected and the entries related to those pathogens were left-out for pathogen-centric zero-shot test. This dataset contained 1,642 unique peptides and 1,861 peptide-pathogen pairs. Complete genome assemblies of the selected 24 species were downloaded from NCBI Genomes (https://www.ncbi.nlm.nih.gov/datasets/genome/). The corresponding accession numbers are provided in Supplementary Table 1.

### Independent In-house Dataset

Using three previously developed *in silico* methods for binary antimicrobial activity prediction^11^, antimicrobial peptide discovery from RNA sequencing datasets^40^, and *de novo* antimicrobial peptide generation^39^, a putative antimicrobial peptide dataset containing 690 peptide sequences was obtained. Consistent with the mined dataset, these peptides did not contain any non-proteinogenic amino acids, were monomeric, and had fewer than 1,022 amino acids. Additionally, these sequences did not comprise any PTMs. To evaluate this independent test set, the peptides were screened against 26 bacterial strains spanning seven species (Supplementary Table 2). Assays were performed according to the protocol of Richter et al^38^, as detailed in the Supplementary Methods.

Raw AST results were first processed where MIC values annotated as “>64” or “>128” μg/mL were converted to 70 and 140 μg/mL, respectively. Next, all MIC measurements were converted from μg/mL to μM. A peptide was labelled positive for a species if it was active (MIC≤32 μM) against any of its strains in any replicate. If it was not active against any strain, it was labelled negative for that species. Consistent with the Mined Dataset, any peptide originally annotated with a “>” whose converted value fell into the positive set was removed. As the last step, the peptide-pathogen pairs that were already included in the Mined Dataset were filtered out. This process yielded a final dataset of 1,354 positive and 1,722 negative peptide-pathogen pairs, spanning 690 peptides and seven species (Supplementary Figure 1). This dataset is termed the in-house dataset.

### Pathogen-Specific Antimicrobial Activity Prediction

#### Embedding Generation

To transform peptide and genome sequences into numerical representations while preserving biologically meaningful features, “esm2_t33_650M_UR50D” variant of ESM2 and “50M_multi_species_v2” variant of NTv2 were employed. For each peptide, ESM2 tokenizes the amino acid sequence, where each token represents a single amino acid residue. Then, for each token, ESM2 generates a numerical vector representation. As a result, for each peptide, the embedding matrix 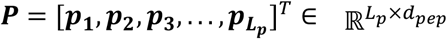 was generated, where ***p***_***i***_ is the embedding vector that represents the *i*^th^ amino acid residue, *L*_*p*_ is the sequence length of the peptide, and *d*_*pep*_ is ESM2’s last layer’s output size (1280).

Following the methodology of NTv2, we converted all the characters other than “A”, “C”, “T”, “G, and “N” to “N” within the 24 genomes in the datasets. These genomes were then broken down into pieces that would each yield 1,000 tokens after merging all the reads for a genome into a single nucleotide sequence (including plasmids). For each resulting piece, an embedding matrix 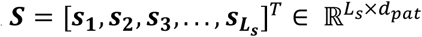 was generated where ***s***_***i***_ is the embedding vector for the *i*^th^ token, *L*_*s*_ is the number of total tokens in the sequence (guaranteed to be 1,000 for all the cases except the last fragment), and *d*_*pat*_ is the output dimension of NTv2’s final layer (512). Next, these 1,000 embedding vectors were aggregated into a single representative vector by mean-pooling. Finally, a genome sequence is represented by the embedding matrix 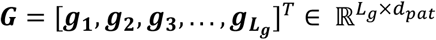, where ***g***_***i***_ is the mean-pooled embedding of the *i*^th^ fragment and *L*_*g*_ is the number of fragments generated from the genome sequence. Finally, embedding matrices of each peptide and pathogen were mean-pooled along their first dimension to obtain per-sequence embeddings.

#### Heterogenous Attention Network-based Link Predictor

For this application, a heterogeneous undirected graph structure *G* = (*V, E, Φ, ψ*) was conceptualized, where *V* represents individual peptides and pathogens, and *E* encompasses peptide-peptide, pathogen-pathogen, and peptide-pathogen links. Function *Φ* maps each node to the categories “peptide” or “pathogen”. Furthermore, function *ψ* maps each edge to the types “peptide similarity” (for peptide-peptide links), “pathogen similarity” (for pathogen-pathogen links), and “bioactivity” (for peptide-pathogen links). To process the associations and extract the hidden patterns in the graph using the outputs of bLLMs, a HLP was proposed.

##### Graph Building

Given a portion of the database containing known positive bioactivity of peptides against pathogens labelled as positive, first, a bipartite graph with “peptide” and “pathogen” node types and “bioactivity” (peptide-pathogen link) edge type was constructed. Then, the bLLM-derived embeddings of the peptides and pathogens are projected into the same feature space with type-specific learnable transformation matrices, to be able to perform message passing, as follows: 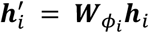, where 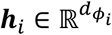 and 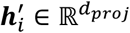 are the original and projected embedding vectors of node 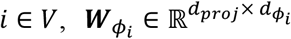 is the type-specific transformation matrix for the node type of node 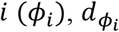 is the size of the original peptide or pathogen embeddings based on the node type of node *i*, and *d*_*proj*_ is the shared size of the projected embedding vectors. Peptide and pathogen similarity edges were then added by calculating the pairwise Euclidean distances between the projected embeddings of nodes of the same type. For each node type, a distance matrix was obtained. Using a specified cutoff percentile (*c*_*pep*_ for peptides or *c*_*pat*_ for pathogens), nodes with a distance smaller than their respective cutoff percentile (excluding self-distances) were connected with their respective “similarity” edges in *G*, ensuring that the embedding projection facilitated the learning of similarities between same-type nodes in terms of antimicrobial activity or susceptibility.

##### Message Passing

The message passing is conducted via metapaths, walks performed on a heterogeneous graph by visiting nodes of a specific order of node types, which define different semantic relationships between the source and destination nodes. For a destination node *i*, the set of source nodes connected to *i* by a composite relation described by the metapath *Φ* is said to be the neighbours of *i* on the metapath *Φ* and symbolized by 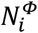. To capture node-level associations and the importance of a node’s neighbours to itself on a given metapath for message passing, Graph Attention Network (GAT)^41^ was employed. On a given metapath *Φ*, the importance of node 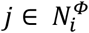 to node *i* for information sharing 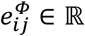 is given by 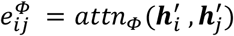, where *attn*_*Φ*_ denotes the self-attention mechanism^42^ for metapath *Φ* and its shared across importance calculations for metapath-*Φ*-based neighbouring node pairs. To use this coefficient as a weight during message passing, the importance of node *j* to node *i* was normalized with the softmax function: 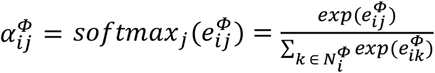, where 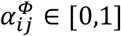 is the weight of node *j* to node *i* on metapath *Φ*, and *exp* is the exponential function. GAT uses a single-layer feedforward neural network as the self-attention function (annotated as *attn*_*Φ*_ above and described in Supplementary Methods). To obtain metapath-*Φ*-based embedding 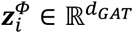 of node *i*, all the messages passed to the node were aggregated using the projected embeddings and attention weights, described as 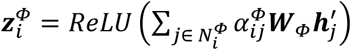 where 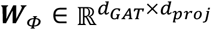 is the linear transformation matrix for the metapath *Φ* and *ReLU* is the Rectified Linear Unit function. Multi-head attention was employed to focus on multiple patterns of the same metapath-based node neighbourships for the final embeddings. The resulting embeddings for each head are concatenated and the final metapath-*Φ*-based embedding 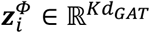 is updated.

##### Semantic Aggregation

As multiple metapaths can be leveraged for node-level attention, multiple embeddings for a single node can be obtained after the message passing. To generate a comprehensive, task-specific representation that summarizes these diverse semantics, we applied a semantic attention module to the metapath-based embeddings. Contrary to the original HAN, we introduced multiple semantic attention modules—one for each node type. These modules learn the importance of metapaths, which are grouped based on their destination node type. Hence, the importance of metapaths targeting a specific node type is normalized only against other metapaths that update that same type.

Given a set of metapaths {*Φ*_1_, *Φ*_2_, . . . *Φ*_*M*_} that ends with the same node type *T*_*V*_′, the node-level attention module generates *M* semantic-specific embeddings for all nodes that have the same node type as the node type of interest, denoted as 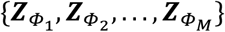. Applied to these sets, semantic attention generates the importances of metapaths as 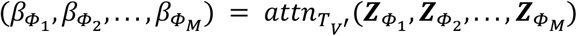 where *attn*_*T*_ ′ is the semantic attention function for the metapaths that updates the node type 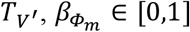 is the weight of metapath *Φ*_*m*_ in the final representation of the nodes and 0 < *m* < *M* + 1 (Supplementary Methods). The final representation of node *i*, 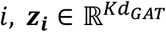, is aggregated as the weighted sum of its metapath-based embeddings with respect to their weights calculated by the semantic attention module as 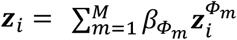.

To enhance the robustness of the semantic attention aggregation, two-dimensional dropout was applied to the metapath-based embeddings. This regularization technique, implemented before the attention calculation, randomly sets a fraction of elements within the semantic-specific embedding vectors to zero during training. This process discourages the model from becoming overly dependent on any single metapath’s representation, forcing it to learn from a more distributed set of features across all available metapaths.

##### Link Prediction

After semantic aggregation of embeddings for both “peptide” and “pathogen” nodes, “bioactivity” link prediction between “peptide” node *i* and “pathogen” node *j* is performed by feeding their final embeddings into a two-layer MLP. This MLP learns the details about the final embeddings of nodes and their interaction with each other in terms of bioactivity. The likelihood of the existence of a link between peptide node *i* and pathogen node *j, p*_*ij*_ ∈ [0, 1], was calculated as *p*_*ij*_ = *sigmoid*(***W***_2_*ReLU*(***W***_1_[***z***_*i*_||*z*_*j*_] + ***b***_1_) + ***b***_2_), where 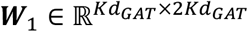 and 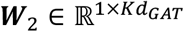 are weight matrices, 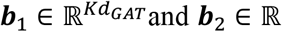 and ***b***_2_ ∈ ℝ are bias terms, and *sigmoid* is the sigmoid function. The interaction between peptide *i* and pathogen *j* is predicted as positive (active) if the corresponding probability score, *p*_*ij*_, exceeds 0.5. Conversely, if *p*_*ij*_ is smaller than or equal to 0.5, the interaction is predicted as negative (inactive).

To train the model using backpropagation of prediction errors, weighted Binary Cross-Entropy Loss (BCEL) was utilized on the true bioactivity status of peptide-pathogen pairs and their corresponding predicted likelihoods for the whole batch as follows:

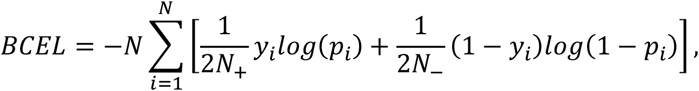

where *y*_*i*_ and *p*_*i*_ are the ground truth bioactivity label and predicted bioactivity probability of the peptide-pathogen pair *i*, respectively, *N* is the number of peptide-pathogen pairs in the supervision set, *N*_+_ and *N*_−_ are the number of positive and negative peptide-pathogen pairs in the supervision set, and *log* is the natural logarithm function. Due to the weighted penalization, the error on the bioactivity class that is less prevalent was emphasized more in the model, acting as a precaution against the class imbalance.

##### Implementation Details

The described HLP was implemented in PyTorch version 2.5.1^43^ and Deep Graph Library (DGL) version 2.4.0^44^, adapting the official DGL HAN baseline implementation (https://github.com/dmlc/dgl/tree/master/examples/pytorch/han). For the hyperparameter tuning and experiments explained in the Results section, projected embedding vector size *d*_*proj*_ was set to 256, the GAT output vector dimension *d*_*GAT*_ was utilized as 64, number of attention heads for GAT *K* was defined as four, the dropout rate for both features and attention weights of GAT were used as 0.5, semantic attention vector size *d*_*s*_ was fixed as 128, and dropout for semantic-specific embeddings for each node was set to 0.5. For optimization of the parameters of the model, the Adam optimizer^45^ with a learning rate of 0.0001 was incorporated, and the model was trained for 2000 epochs. Each step of the optimizer was taken after a full iteration on the supervision edges. Six distinct directed (or four undirected) metapaths representing four fundamental interaction categories, were selected for the system. The rationale behind these metapaths was to extract latent information regarding: (1) peptide similarity based on shared pathogen targets (“peptide” to “pathogen” to “peptide” metapath); (2) pathogen similarity based on susceptibility to common peptides (“pathogen” to “peptide” to “pathogen” metapath); (3) potential peptide activity inferred from similar peptides active against a specific pathogen (“peptide” to “peptide” to “pathogen” metapath and its reverse); and (4) pathogen susceptibility inferred from similar pathogens being susceptible to a specific peptide (“pathogen” to “pathogen” to “peptide” metapath and its reverse). Three hyperparameters were optimized: similarity cutoffs for graph building and the mechanism for incorporating self-embeddings of nodes. For the similarity cutoffs, {5, 10, 15} for peptides and {10, 20, 30} for pathogens. Concurrently, the search compared two distinct methods for integrating self-embeddings: adding them as self-loops within standard metapaths, allowing node-level attention to dynamically weigh the self-embedding against its neighbors, versus treating them as an independent conceptual metapath where their overall contribution is collectively determined by a single, shared weight learned by the semantic attention module. For the selection of both the best weights for a hyperparameter combination and the best hyperparameter configuration, macro F1-score (the average of pathogen-specific F1-Scores) was used (Supplementary Tables 4 and 5).

#### Conventional Machine Learning Approaches

##### Multilayer Perceptron

A deeper MLP that classifies the bioactivity of peptide *i* against pathogen *j* was trained on the bLLM-derived embeddings. A single perceptron of the MLP classifier was defined as 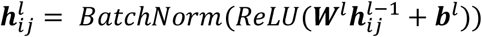 where 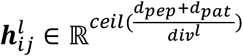 is the output vector of the *l*^th^ perceptron, 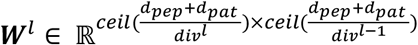 is the weight matrix of the *l*^th^ perceptron, 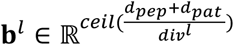 is the bias vector of the *l*^th^ perceptron, *ReLU* is the Rectified Linear Unit function, *BatchNorm* is the batch normalization function, *ceil* is the ceiling function and *div* is the shrinking factor for each perceptron stacked, defined as a hyperparameter. Here, *l* starts from 1 and ends at where the output size of the perceptron is one. In case of *l*=1, the embeddings of peptide *i* and pathogen *j* were concatenated as the input (i.e. 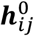), and when the output dimension was one, the batch normalization and *ReLU* activation were replaced with the sigmoid function. The input vector for the first perceptron, 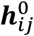, was defined as 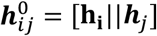, where 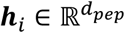 represents the BLLM-based embedding of peptide 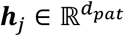 represents the BLLM-based embedding of pathogen *j, d*_*pep*_ and *d*_*pat*_ denote the size of protein and pathogen embeddings, respectively, and || is the concatenation operation. Error was calculated using a weighted BCEL, where positive samples were scaled by the ratio of negative-to-positive instances to balance the contribution of each class. The characterized MLP was implemented using PyTorch version 2.5.1 and trained for 50 epochs. The AdamW optimizer^46^ was used to optimize the parameters. The hyperparameter shrinking factor *div* was tuned by testing the discrete values {2, 4, 8, 16, 32, 64} based on macro F1-scores (Supplementary Table 6).

##### SVM

Consistent with the MLP model, feature vectors for each peptide-pathogen pair were generated by concatenating their respective embeddings. An SVM binary classifier, implemented in Scikit-learn version 1.5.2, was then used to predict the targeted bioactivity. To ensure consistency with other models in the ensemble, SVM prediction probabilities were utilized rather than direct class predictions. However, as noted in the Scikit-learn documentation (https://scikit-learn.org/stable/modules/generated/sklearn.svm.SVC.html), prediction probabilities may occasionally diverge from direct predictions. This discrepancy is acknowledged as a limitation of the study. The SVM kernel was treated as a hyperparameter, and the validation Macro F1-Score was optimized by testing the set {linear, third-degree polynomial, radial basis function}. Every other parameter was set to its default value (Supplementary Table 7).

##### XGBoost

Adopting the same feature representation as the MLP and SVM models, concatenated peptide-pathogen embeddings were used as input for an XGBoost binary classifier to predict bioactivity. The classifier was implemented using the XGBoost Python library version 2.1.4 (https://xgboost.readthedocs.io/en/stable/index.html). The model was trained with a learning rate of 0.01, using binary logistic regression as the objective function. A hyperparameter search was conducted to optimize the Macro F1-Score on the validation set. The search space included the number of estimators:

{200, 400, 600, 800, 1000, 1200, 1400, 1600, 1800, 2000}, and maximum depth of the trees: {2, 4, 6, 8, 10, 12, 14, 16, 18, 20}. The combination of the number of estimators and maximum depth of the trees that returned the highest validation Macro F1-Score was selected for the final mode (Supplementary Table 8).

#### Blending Ensemble

To obtain a final predictor that exploits the predictions of the classifiers discussed above, a Logistic Regression meta learner was trained on the validation dataset, as a Blending strategy, using the output logits of the classifiers as features. The selection of Logistic Regression was made due to its simplicity and effectiveness on non-complex data, as the logit outputs of up to four models were given to the model for binary classification. It was implemented using the Scikit-learn version 1.5.2. L2 regularization was applied with a regularization coefficient of 100 as a precaution against overfitting on smaller data, like the validation set. Other parameters of the model were left at their default.

### Data Split Schemes

We developed two graph learning schemes that redefine data partitioning. In both schemes, each stage of development requires two sets of edges: a message passing set, where the information flows through, and a supervision set, where the model performs link prediction and the performance is measured on.

Note that the “peptide similarity” and “pathogen similarity” edges were constructed dynamically based on the outputs of the node-type-specific projection matrices, after the initial graph was built with “bioactivity” edges. Hence, they were not partitioned; rather, they were built based on the given graph with “bioactivity” edges established.

### Transductive Learning

Following the HLP notation, we partitioned the database by splitting the “bioactivity” edges, while keeping all “peptide” and “pathogen” nodes present in each graph. First, all known positive and negative peptide-pathogen pairs (edges) were partitioned into training (70%), validation (10%), and test (20%) sets. During the training phase, the training set was split: 50% of its positive edges were used for message passing, while the other 50% of positive edges and all its negative edges were used for supervision. For validation, the graph structure was expanded: all positive edges from the training set (70%) were used for message passing, and the model was evaluated using the validation set (10%). Similarly, for testing, the message passing graph included all positive edges from both the training and validation sets (80% combined), and the model was evaluated on the testing set (20%). This strategy was called Transductive Learning, as no new nodes were introduced during validation or testing. Including all nodes at all stages enables the model to learn the complete distribution of peptide embeddings and their inter-relationships. The trade-off of this transductive design is reduced generalizability, as the model is never validated or tested on new peptides.

### Inductive Learning

For the Inductive Learning scheme, “peptide” nodes were partitioned into three sets, while all the “pathogen” nodes were included in all the graphs due to their small number. Here, for a given split, Training, Validation, and Testing, all the “bioactivity” edges connected to the corresponding “peptide” nodes were partitioned into the supervision set, and 50% of those edges were used for message passing. In contrast, the other half was used for supervision and/or evaluation. The primary advantage of this inductive approach is its ability to generalize to new nodes. However, this comes at the cost of losing global context, as the model is trained on a smaller set and cannot see the entire dataset’s structure at once.

To ensure a fair comparison, the same supervision edges used for the HLP were also used to train and evaluate three conventional machine learning methods. Specifically, the supervision edge sets from the training, validation, and test splits were directly reused for the training and inference stages of the conventional machine learning models.

## CODE AND DATA AVAILABILITY

All source code, model weights, and the mined dataset used in this study are publicly accessible via GitHub at https://github.com/BirolLab/triAMPh. The in-house dataset is available from the corresponding author upon reasonable request.

## GENERATIVE ARTIFICIAL INTELLIGENCE USE

During the preparation of this work, the authors utilized generative artificial intelligence tools (specifically ChatGPT 4.0 and Gemini 3 Pro) to assist with text editing and code review. The authors take full responsibility for all scientific content, data analysis, and interpretations presented in this manuscript.

## CONFLICT OF INTEREST

Amphoraxe Life Sciences Inc. has licensed the intellectual property (IP) associated with some of the antimicrobial peptides contained in the in-house dataset. I.B., who co-invented this IP alongside A.Y., V.C.T, R.L.W., and C.C.H., also serves as a founder and Chief Scientific Officer of Amphoraxe and holds equity in the company. Additionally, E.D., A.S., and B.U. were supported by Amphoraxe-backed Mitacs internships. All authors confirm that the company was entirely excluded from data collection, data analysis, results interpretation, and the decision to submit for publication. All other authors declare no competing interests.

## FUNDING

This work was financially supported by Genome Canada and Genome British Columbia through the Large Scale Applied Projects program (#291PEP) awarded to I.B. and C.C.B. Additionally, trainees received partial support from Mitacs (IT28945, IT43363), the University of British Columbia, and Amphoraxe Life Sciences Inc.

## Notes

https://github.com/BirolLab/triAMPh

