## Supplementary Material for "Pathogen-specific antimicrobial activity prediction with biological large language model-based methods"

### SUPPLEMENTARY METHODS

#### In-house Database Antimicrobial Susceptibility Testing

A total of 26 bacterial strains, comprising seven species, were obtained from two sources: purchased from the American Type Culture Collection (ATCC; Manassas, VA, USA) or as human and bird clinical isolates by the British Columbia Centre for Disease Control and Vaccine and Infectious Disease Organisation-InterVac (VIDO-InterVac; Saskatoon, SK, CA). The detailed information about the strains can be found in Table A.2. Bacteria were cultured overnight at 37 °C in a shaking incubator using Mueller-Hinton Broth (MHB; Sigma-Aldrich, St. Louis, MO, USA). No additives were included in the MHB during cultivation. For storage, the overnight cultures were mixed in a 1:1 ratio with a 50% glycerol solution, dispensed into cryovials, and stored at −80 °C.

Bacterial glycerol stocks from −80 °C storage were streaked onto Columbia blood agar (MP0351; Oxoid, Nepean, ON, Canada) and incubated overnight at 37 °C. To ensure a healthy and uniform population for the assay, 2–4 colonies were subcultured onto a fresh plate. After incubation, isolated colonies were suspended in MHB. This suspension was standardized using a spectrophotometer to an initial optical density corresponding to $\approx 1\times10^{8}$ colony-forming unit per millilitre (CFU/mL), which corresponds to 0.5 McFarland standard. This stock was then diluted 1:250 to create the final working inoculum of $5 \pm 3 \times10^{5}$ CFU/mL. The concentration, purity, and viability of this inoculum were confirmed for each trial using a Total Viability Count, which involved spread-plating a 1:1,000 dilution on nonselective media with duplicates.

For the antimicrobial susceptibility testing (AST) assays, first, UltraPure water (Thermo Fisher Scientific, Waltham, MA, USA) was used to resuspend lyophilized 80 μg peptide aliquots (GenScript; Piscataway, NJ, USA) to a 1.28 mg/mL stock solution. These peptides were serially diluted to establish a concentration gradient from 128 μg/mL down to 0.25 μg/mL or 64 μg/mL to 0.125 μg/mL, in 96-well polypropylene microtiter plates (Greiner Bio-One #650261, Kremsmünster, Austria). Growth and sterility controls (MHB with bacteria and only MHB, respectively) were represented in two columns per plate. The prepared bacterial inoculum was dispensed into all peptide-containing wells and the growth control wells. Following a 20–24 hour incubation at 37 °C, for the ASTs done manually the minimum inhibitory concentration (MIC) was determined by visual inspection as the lowest peptide concentration that inhibited any visible growth.

#### Database Partitioning for Competitor Benchmarking

For AMPSpeciesSpecific[1], the model and prediction code were downloaded from its GitHub repository (https://github.com/bzlee-bio/AMPSpeciesSpecific). This model predicts peptide activity for five specific species: *Bacillus subtilis, Escherichia coli, Pseudomonas aeruginosa, Staphylococcus aureus*, and *Staphylococcus epidermidis*. The mined dataset's test subset contains pairs for all five of these species, while the in-house set encompasses three of them (*E. coli, P. aeruginosa, and S. aureus*). Both datasets were subsequently filtered to include only the peptide-pathogen pairs associated with these shared target species. Using default parameters, the tool's inference file was then executed on these reduced datasets. The AMPSpeciesSpecific model bases its classification on a 267.7 µg/mL activity threshold (averaged MIC). Its authors also used UniProt[2] peptides (resulting from a "NOT antimicrobial" search) as their negative set. As this study exclusively uses AMP databases to exploit known efficacy measurements, this threshold was considered too generous as most of the measurements done In-house did not go over 256 µg/mL. Therefore, no additional subsetting of our test sets was performed based on this activity threshold. Another important pre-processing difference was the handling of duplicate pairs. AMPSpeciesSpecific calculates the mean MIC for identical peptide-pathogen pairs. In this study, however, a single active measurement (MIC ≤ 32 µM) was deemed sufficient to classify a pair as positive. This methodological discrepancy was also not accounted for during data subsetting.

The model developed by Vishnepolsky et al.[2] provides a web server (https://dbaasp.org/tools?page=linear-amp-prediction) for its strain-specific peptide bioactivity classifier, which supports eight trained strains (*Acinetobacter baumannii* ATCC 19606, *Bacillus subtilis* ATCC 6633, *E. coli* ATCC 25922, *Enterococcus faecalis* ATCC 29212, *Klebsiella. pneumoniae* ATCC 700603, *P. aeruginosa* ATCC 27853, *S. aureus* ATCC

25923, and *Salmonella typhimurium* ATCC 14028) and allows user-uploaded genomes. This baseline presented two major challenges for comparison. First, our study uses species-level reference genomes, which are not an accurate match for a strain-specific classifier. Second, the authors define bioactivity using different thresholds: active if MIC < 25 µg/ml and inactive if MIC > 100 µg/ml.

To create a valid test subset, we modified our dataset-building process to select only peptide-species pairs with mutually consistent activity labels across both studies. For positive pairs: A pair was included only if it was active in our dataset (i.e., had at least one measurement with MIC ≤ 32 µM against any strain of that species), and at least one of those measurements contained the ATCC identification number of a specific strain provided by Vishnepolsky et al. and also met their active threshold (MIC < 25 µg/mL). For negative pairs: A pair was included only if it was classified as negative in our dataset (meaning no positive measurements ≤ 32 µM existed), and at least one of its negative measurements was against a Vishnepolsky et al. strain and also met their inactive threshold (MIC > 100 µg/mL). After applying this filtering, the mined dataset test subset included pairs for all eight strains. The in-house Dataset subset contained pairs for four of the strains (*A. baumannii* ATCC 19606, *E. coli* ATCC 25922, *E. faecalis* ATCC 29212, and *P. aeruginosa* ATCC 27853).

A further discrepancy was the Vishnepolsky et al. model's ability to process C-terminal amidation, a modification that our study disregards. To ensure a fair comparison and align the datasets, we implemented a specific rule during our pre-processing, prior to eliminating duplicates. When examining a peptide's C-terminal modification status, if it was found in both modified (amidated) and non-modified forms, the non-modified version was prioritized. This was done for greater concordance with our own method. However, if a peptide was only available in an amidated form, a "+" was added to its sequence—as required by their model—to allow for its correct processing. Moreover, peptides with unknown C-terminal modifications were assumed to be non-modified. This was done as exclusion of those pairs would shrink the dataset substantially. Additionally, due to Vishnepolsky et al.'s data collation, the peptides, we excluded peptides shorter than 9 or longer than 30 amino acids as well as peptides containing N-terminal modifications or unknown status for it. Here, we observed some of the data pairs being annotated with wrong ATCC identifiers. We removed pairs which were annotated with *Enterococcus faecium* instead of *E. faecalis* for ATCC 29212.

When we compared our final mined test set subset to training set of Vishnepolsky et al.'s model, we observed seven of the mined test set pairs annotated with the incorrect label compared to the public databases at the time of this study. This might be due to additional data being pushed to these databases after they concluded their study.

#### Heterogenous Attention Network-based Link Predictor

##### Message Passing

The final attention weights obtained from the Graph Attention Network (GAT)[3] can be expressed in detail as $\alpha_{ij}^{\Phi} = \frac{exp(LeakyReLU(\boldsymbol{a}_{\Phi}^{T}\,[\boldsymbol{W}_{\Phi}\boldsymbol{h}_i' || \boldsymbol{W}_{\Phi}\boldsymbol{h}_j']))}{\sum_{k\,\in\,N_i^{\Phi}} exp(LeakyReLU(\boldsymbol{a}_{\Phi}^{T}\,[\boldsymbol{W}_{\Phi}\boldsymbol{h}_i' || \boldsymbol{W}_{\Phi}\boldsymbol{h}_k']))}$, where $\boldsymbol{W}_{\Phi} \in \mathbb{R}^{d_{GAT} \times d_{proj}}$ is the linear transformation matrix for the metapath $\Phi$, $\boldsymbol{a}_{\Phi} \in \mathbb{R}^{2d_{GAT}}$ is the attention vector calculating how much weight should be given to the current pair of nodes based on their projected embeddings for the metapath $\Phi$, $d_{GAT}$ is the output size of a single GAT head, $LeakyReLU$ is the Leaky Rectified Linear Unit function ($LeakyReLU(x) = max(0, x) + 0.1 \times min(0, x)$) for the addition of nonlinearity, and $||$ indicates the concatenation operation.

To enhance the robustness of the node-level attention mechanism, a dual-dropout strategy was implemented. First, dropout is applied to the node feature matrices of the neighbourhood before the attention computation. Second, dropout is applied directly to the unnormalized attention coefficients before they are passed through the softmax normalization function. This approach prevents the model from over-relying on specific neighbours and compels it to learn a more distributed attention weighting across a stochastically sampled neighbourhood.

#### Semantic Aggregation

To calculate the weight of a metapath $\Phi_m$, first, each of the metapath-$\Phi_m$-based embeddings was subjected to a nonlinear transformation via an MLP. Consequently, the average similarity of the outputs of this MLP for all the embeddings on metapath $\Phi_m$ to a learnable attention vector $\boldsymbol{q} \in \mathbb{R}^{d_s}$ is assigned as the importance $w_{\Phi_m} \in \mathbb{R}$. Semantic attention steps can be represented in detail as $w_{\Phi_m} = \frac{1}{|V_{T_{V'}}|}\sum_{i \in V_{T_{V'}}} \boldsymbol{q}^T tanh(\boldsymbol{W}\boldsymbol{z}_i^{\Phi_m} + \boldsymbol{b})$, where $\boldsymbol{W} \in \mathbb{R}^{d_s \times K d_{GAT}}$ is the weight matrix, $\boldsymbol{b} \in \mathbb{R}^{d_s}$ is the bias vector, $tanh$ is the activation function, $V_{T_{V'}}$ is the set of nodes that have the note type $T_{V'}$, and $|\cdot|$ is the cardinality operation. To normalize the importance of the metapath $\Phi_m$ and obtain its weight, the softmax function was used as $\beta_{\Phi_m} = \frac{exp(w_{\Phi_m})}{\sum_{j=1}^{M} exp(w_{\Phi_j})}$.

### SUPPLEMENTARY FIGURES

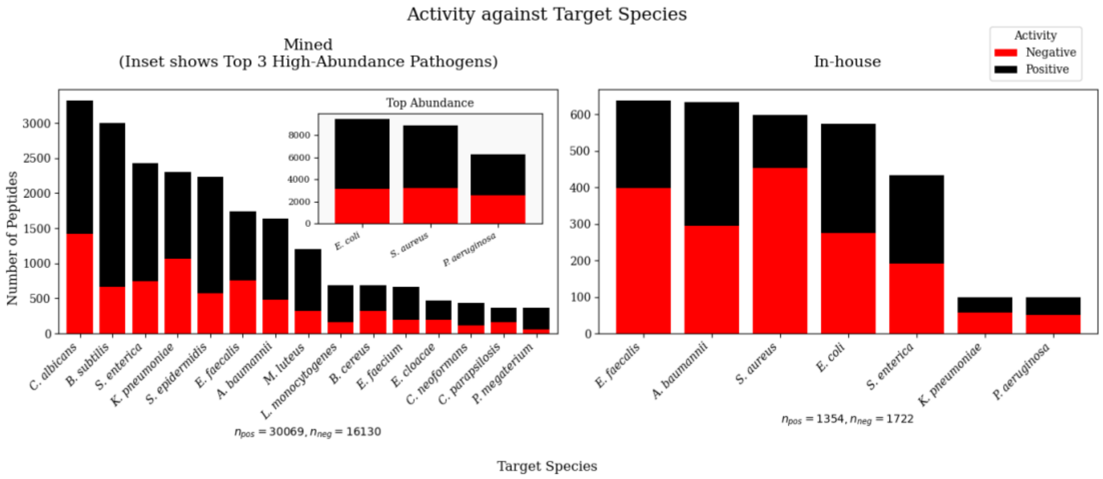

**Supplementary Figure 1. Distribution of peptide-pathogen pairs stratified by target species for the Mined (Left) and In-house (Right) Datasets.** Stacked bars represent the number of active (Black) and inactive (Red) peptides experimentally validated against each species. To resolve scale differences, the three most frequently tested species are displayed in an inset panel, while the remaining species are plotted on the main axes.

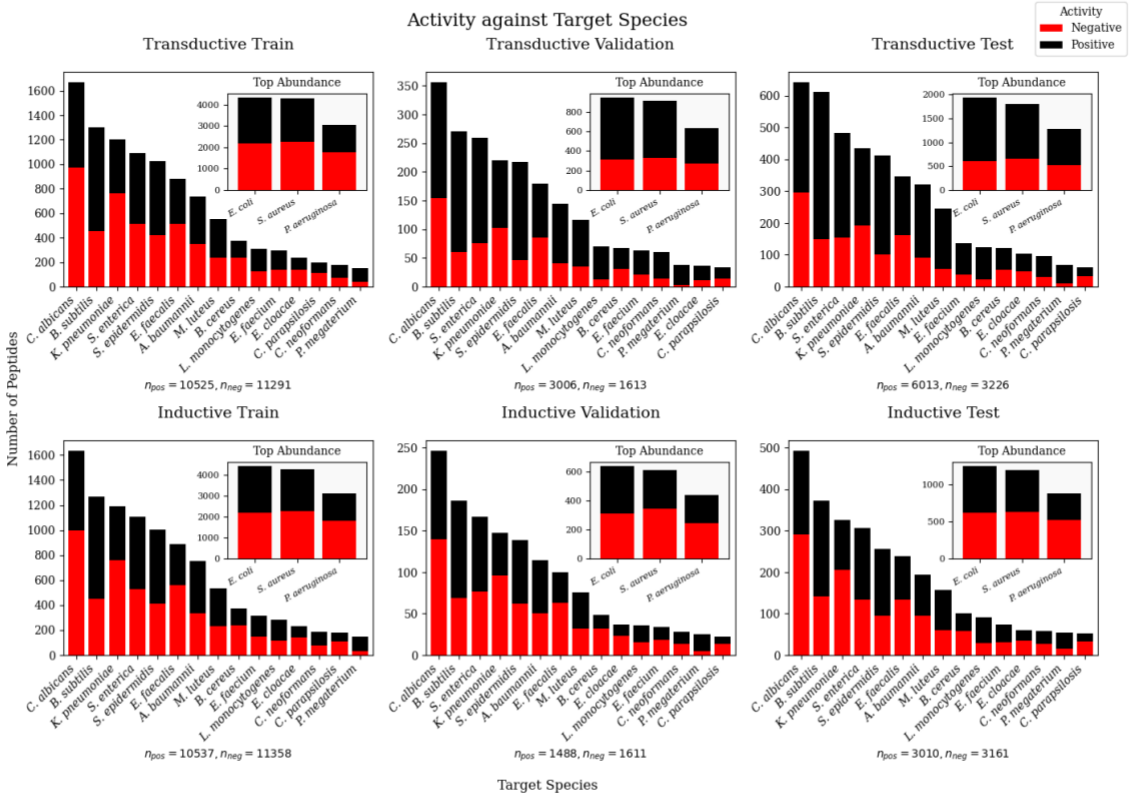

**Supplementary Figure 2. Distribution of peptide-pathogen pairs across target species for the Transductive (Top row) and Inductive (Bottom row) supervision splits.** Stacked bars represent the number of active (Black) and inactive (Red) peptides experimentally validated against each species. Insets display the most frequently tested species to accommodate the substantial scale disparity and preserve the readability of the remaining data

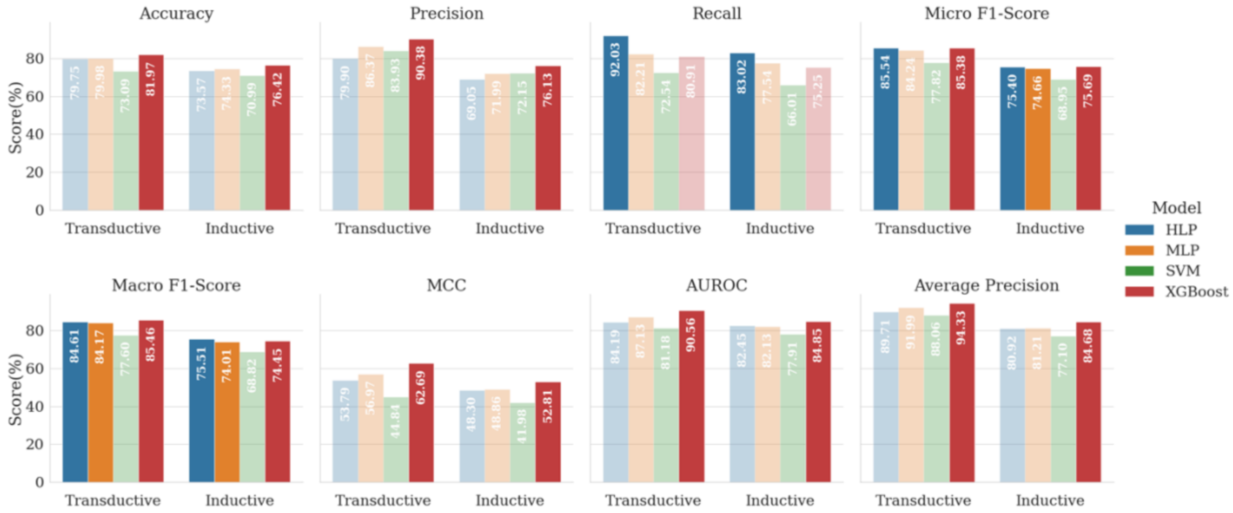

**Supplementary Figure 3. Performance of individual machine learning models on the mined test set, across eight metrics.** The data tested contained 9,239 and 6,171 bioactivity pairs for Transductive and Inductive Learning schemes, respectively. The transparency denotes statistical significance (p-value < 0.01) in performance gap compared to the best-performing model in the same learning scheme based on 100,000 bootstrapping iterations followed by Holm-Bonferroni correction.

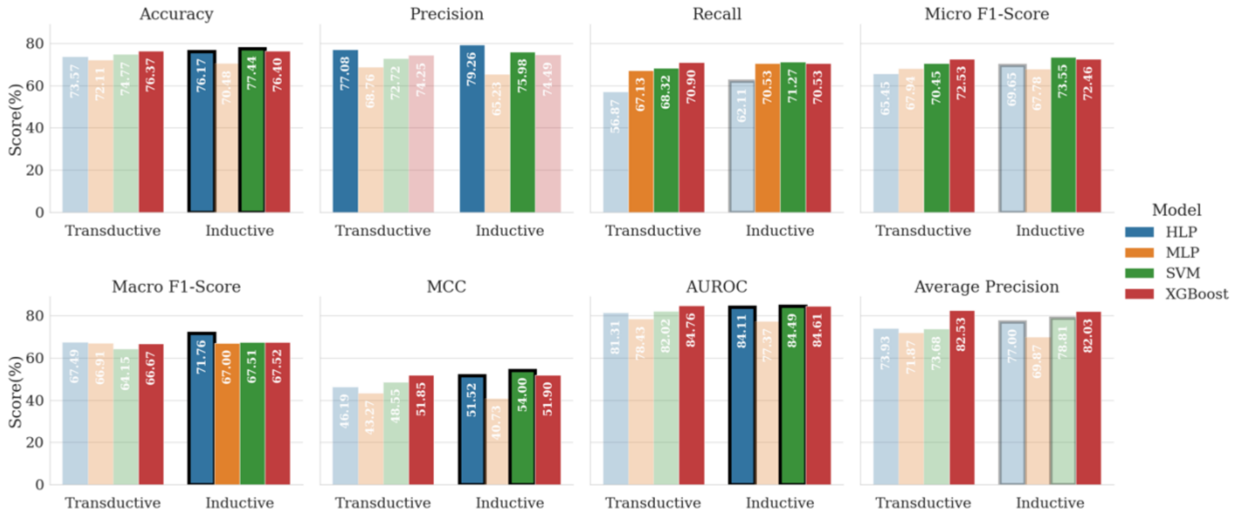

**Supplementary Figure 4. Performance of individual machine learning models on the in-house set, across eight metrics.** The dataset contained 3,076 bioactivity pairs. Transparency denotes statistical significance (p-value < 0.01) in performance gap compared to the best-performing model for the same metric based on 100,000 bootstrapping iterations followed by Holm-Bonferroni correction. Black borders denote statistical significance (p<0.01) for the performance gap relative to the same model in the alternative learning scheme.

### SUPPLEMENTARY TABLES

**Supplementary Table 1. Regularization rules applied to the AMP databases' reported MICs for peptide-pathogen pairs.** Letters x and y denote numbers.

| Original Occurrence | Converted Value |
|---|---|
| $> x$ | $2x$ |
| $\geq x$ | $x\sqrt{2}$ |
| $x$ - $y$ | $\sqrt{xy}$ |
| $x$ -$\geq y$ | $\sqrt{xy\sqrt{2}}$ |
| $< x$ | $x$ |
| $\leq x$ | $x$ |
| $x + y$ | $\sqrt{x(x+y)}$ |
| $x \pm y$ | $\sqrt{(x-y)(x+y)}$ |

**Supplementary Table 2. The species included in the mined dataset and the respective genome assemblies used for embedding generation.** The source of the genome assemblies was noted in parentheses.

| Species | Accession |
|---|---|
| *Acinetobacter baumannii* | GCF_009035845.1 (NCBI RefSeq) |
| *Bacillus cereus* | GCF_002220285.1 (NCBI RefSeq) |

| Species | Accession |
| --- | --- |
| *Bacillus subtilis* | GCF_000009045.1 (NCBI RefSeq) |
| *Candida albicans* | GCF_000182965.3 (NCBI RefSeq) |
| *Candida parapsilosis* | GCA_000182765.2 (GenBank) |
| *Candida tropicalis* | GCF_000006335.3 (NCBI RefSeq) |
| *Cryptococcus neoformans* | GCF_000091045.1 (NCBI RefSeq) |
| *Enterobacter cloacae* | GCF_905331265.2 (NCBI RefSeq) |
| *Enterococcus faecalis* | GCA_000393015.1 (GenBank) |
| *Enterococcus faecium* | GCF_009734005.1 (NCBI RefSeq) |
| *Escherichia coli* | GCF_000005845.2 (NCBI RefSeq) |
| *Klebsiella pneumoniae* | GCF_000240185.1 (NCBI RefSeq) |
| *Listeria monocytogenes* | GCF_000196035.1 (NCBI RefSeq) |
| *Micrococcus luteus* | GCF_900475555.1 (NCBI RefSeq) |
| *Priestia megaterium* | GCF_006094495.1 (NCBI RefSeq) |
| *Proteus mirabilis* | GCF_000069965.1(NCBI RefSeq) |
| *Pseudomonas aeruginosa* | GCF_000006765.1 (NCBI RefSeq) |
| *Pseudomonas syringae* | GCF_018394375.1(NCBI RefSeq) |
| *Salmonella enterica* | GCF_000006945.2 (NCBI RefSeq) |
| *Saccharomyces cerevisiae* | GCF_000146045.2 (NCBI RefSeq) |
| *Staphylococcus aureus* | GCF_000013425.1 (NCBI RefSeq) |
| *Staphylococcus epidermidis* | GCF_006094375.1 (NCBI RefSeq) |
| *Streptococcus mutans* | GCF_019048645.1 (NCBI RefSeq) |
| *Streptococcus pyogenes* | GCF_900475035.1 (NCBI RefSeq) |

**Supplementary Table 3. The details of the strains used in ASTs done in-house.**

| **Strains Tested** |
|---|
| *Acinetobacter baumannii* (strain 2208) ATCC 19606 |
| *Acinetobacter baumannii* NDM/OXA-51 (BCCDC clinical isolate) |
| *Acinetobacter baumannii* ATCC 1710 |
| *Enterococcus faecalis* ATCC 29212 |
| *Enterococcus faecalis* vanA (BCCDC clinical isolate) |
| *Escherichia coli* ATCC 25922 |
| Avian Pathogenic *Escherichia coli* 317 |
| *Escherichia coli* ATCC BAA-2469 |
| *Escherichia coli* ATCC BAA-2340 |
| *Escherichia coli* ATCC BAA-2471 |
| *Escherichia coli* CPO-NDM (BCCDC clinical isolate) |
| *Escherichia coli* CPO-KPC (BCCDC clinical isolate) |
| *Escherichia coli* ESBL (BCCDC clinical isolate) |
| *Escherichia coli* MCR-1 (BCCDC clinical isolate) |
| *Escherichia coli* MCR-2 (BCCDC clinical isolate) |
| *Escherichia coli* MCR/NDM (BCCDC clinical isolate) |
| *Klebsiella pneumoniae* ATCC 10031 |
| *Klebsiella pneumoniae* KPC/NDM (BCCDC clinical isolate) |
| *Pseudomonas aeruginosa* ATCC 27853 |
| *Pseudomonas aeruginosa* NDM/VIM (BCCDC clinical isolate) |
| *Salmonella enterica* serovar *Enteritidis* ATCC 4931 |
| *Salmonella enterica* serovar Enteritidis LS101 |
| *Salmonella enterica* serovar Enteritidis (BCCDC clinical isolate) |

| **Strains Tested** |
|---|
| *Salmonella enterica* serovar Heidelberg (BCCDC clinical isolate) |
| *Staphylococcus aureus* ATCC 29213 |
| *Staphylococcus aureus* (BCCDC Clinical Isolate) |

**Supplementary Table 4. Hyperparameter tuning results of HLP for the Transductive Learning scheme.** Supervision Validation portion of the Transductive splits which contains 4,619 pairs was used. All values are reported as percentages. The best-performing metric values are highlighted in bold.

| **Self-Embedding Inclusion** | **Peptide Cutoff** | **Pathogen Cutoff** | **Accuracy** | **Precision** | **Recall** | **Micro F1-Score** | **Macro F1-Score** | **MCC** | **AUROC** | **Average Precision** |
|---|---|---|---|---|---|---|---|---|---|---|
| Node-level | 5 | 10 | **79.45** | **80.19** | 90.88 | **85.20** | **85.26** | **53.14** | 83.45 | 89.20 |
| | | 20 | 78.74 | 78.38 | 92.98 | 85.06 | 85.01 | 51.34 | 83.32 | 89.01 |
| | | 30 | 78.78 | 78.46 | 92.91 | 85.07 | 85.11 | 51.44 | 83.40 | 89.00 |
| | 10 | 10 | 78.93 | 78.76 | 92.61 | 85.12 | 85.03 | 51.81 | **83.58** | **89.28** |
| | | 20 | 78.68 | 78.23 | 93.15 | 85.04 | 84.93 | 51.19 | 83.23 | 88.93 |
| | | 30 | 78.83 | 78.76 | 92.38 | 85.03 | 84.88 | 51.54 | 83.21 | 88.83 |
| | 15 | 10 | 79.04 | 79.13 | 92.08 | 85.12 | 85.15 | 52.07 | 83.19 | 88.93 |
| | | 20 | 78.72 | 78.29 | 93.11 | 85.06 | 85.04 | 51.30 | 83.13 | 88.83 |
| | | 30 | 78.50 | 77.84 | **93.61** | 85.00 | 85.13 | 50.80 | 83.40 | 88.95 |
| Semantic-level | 5 | 10 | 77.40 | 78.34 | 90.22 | 83.86 | 84.05 | 48.10 | 80.79 | 86.83 |
| | | 20 | 77.42 | 78.46 | 90.02 | 83.84 | 83.87 | 48.18 | 81.33 | 87.19 |
| | | 30 | 77.38 | 78.18 | 90.49 | 83.89 | 83.92 | 48.02 | 80.70 | 86.73 |
| | 10 | 10 | 77.42 | 78.36 | 90.22 | 83.87 | 84.05 | 48.16 | 80.78 | 86.83 |
| | | 20 | 77.35 | 79.08 | 88.66 | 83.59 | 83.88 | 48.24 | 81.03 | 86.91 |
| | | 30 | 77.38 | 78.18 | 90.49 | 83.89 | 83.90 | 48.02 | 80.70 | 86.73 |

| Self-Embedding Inclusion | Peptide Cutoff | Pathogen Cutoff | Accuracy | Precision | Recall | Micro F1-Score | Macro F1-Score | MCC | AUROC | Average Precision |
|---|---|---|---|---|---|---|---|---|---|---|
| | 15 | 10 | 77.38 | 78.38 | 90.09 | 83.83 | 84.03 | 48.07 | 80.78 | 86.83 |
| | | 20 | 77.38 | 79.07 | 88.72 | 83.62 | 83.89 | 48.28 | 81.03 | 86.92 |
| | | 30 | 77.33 | 78.14 | 90.49 | 83.86 | 83.89 | 47.91 | 80.70 | 86.73 |

**Supplementary Table 5. Hyperparameter tuning results of HLP for the Inductive Learning scheme**. Supervision Validation portion of the Inductive splits which contains 3,099 pairs was used. All values are reported as percentages. The best-performing metric values are highlighted in bold.

| Self-Embedding Inclusion | Peptide Cutoff | Pathogen Cutoff | Accuracy | Precision | Recall | Micro F1-Score | Macro F1-Score | MCC | AUROC | Average Precision |
|---|---|---|---|---|---|---|---|---|---|---|
| Node-level | 5 | 10 | 72.09 | 65.90 | 86.76 | 74.91 | 74.31 | 46.93 | 76.06 | 69.39 |
| | | 20 | 71.83 | 65.52 | 87.23 | 74.83 | 74.28 | 46.64 | 76.38 | 69.63 |
| | | 30 | 71.77 | 65.44 | 87.30 | 74.81 | 74.23 | 46.56 | 76.72 | 70.29 |
| | 10 | 10 | 73.06 | 66.84 | 87.10 | 75.63 | 74.24 | 48.71 | 80.35 | 77.14 |
| | | 20 | 71.41 | 65.00 | 87.63 | 74.64 | 74.10 | 46.07 | 75.71 | 68.65 |
| | | 30 | 73.15 | 66.82 | 87.57 | 75.80 | 74.24 | 49.03 | 80.22 | 75.95 |
| | 15 | 10 | 71.99 | 65.72 | 87.10 | 74.91 | 74.18 | 46.88 | 75.86 | 68.64 |
| | | 20 | 72.35 | 65.91 | 87.84 | 75.31 | 74.20 | 47.74 | 80.25 | 75.96 |
| | | 30 | 73.22 | 67.19 | 86.42 | 75.60 | **74.40** | 48.77 | 80.67 | 76.84 |
| Semantic-level | 5 | 10 | 73.25 | 67.01 | 87.23 | 75.80 | 74.19 | 49.09 | **81.86** | **79.08** |
| | | 20 | 72.93 | 66.20 | **89.11** | **75.97** | 74.16 | **49.21** | 81.67 | 78.90 |
| | | 30 | 73.41 | 67.57 | 85.82 | 75.61 | 74.14 | 48.93 | 81.71 | 78.97 |
| | 10 | 10 | 73.41 | 67.60 | 85.69 | 75.58 | 74.08 | 48.89 | 81.62 | 78.78 |
| | | 20 | 73.41 | 67.38 | 86.49 | 75.75 | 74.05 | 49.13 | 81.24 | 78.21 |

| Self-Embedding Inclusion | Peptide Cutoff | Pathogen Cutoff | Accuracy | Precision | Recall | Micro F1-Score | Macro F1-Score | MCC | AUROC | Average Precision |
|---|---|---|---|---|---|---|---|---|---|---|
| | | 30 | 73.83 | **68.80** | 83.27 | 75.34 | 74.13 | 48.99 | 81.57 | 78.82 |
| | 15 | 10 | 72.86 | 66.62 | 87.16 | 75.52 | 74.15 | 48.40 | 81.62 | 78.78 |
| | | 20 | 73.35 | 67.49 | 85.82 | 75.56 | 74.05 | 48.81 | 81.54 | 78.62 |
| | | 30 | **73.89** | 68.77 | 83.60 | 75.46 | 74.19 | 49.19 | 81.55 | 78.82 |

**Supplementary Table 6. Hyperparameter tuning results for the MLP on the respective supervision Validation set**. The sizes of the Transductive and Inductive dataset partitions used were 4,619 and 3,099 bioactivity pairs, respectively. All values are reported as percentages. Divisor is the divisor used to divide the size of the current perceptron by to obtain the size of the next perceptron.

| Learning | Divisor | Accuracy | Precision | Recall | Micro F1-Score | Macro F1-Score | MCC | AUROC | Average Precision |
|---|---|---|---|---|---|---|---|---|---|
| Transductive | 2 | 80.08 | 86.02 | 82.87 | 84.41 | 83.73 | 56.94 | 86.34 | 91.36 |
| | 4 | 78.74 | 85.94 | 80.51 | 83.13 | 83.40 | 54.68 | 85.49 | 90.80 |
| | 8 | 77.46 | 87.37 | 76.41 | 81.53 | 81.98 | 53.75 | 85.42 | 91.06 |
| | 16 | 78.37 | 84.21 | 82.17 | 83.18 | 82.17 | 52.94 | 84.54 | 90.38 |
| | 32 | 74.09 | 78.83 | 82.27 | 80.51 | 80.40 | 41.98 | 79.40 | 86.49 |
| | 64 | 73.48 | 78.92 | 80.84 | 79.87 | 79.52 | 41.06 | 78.43 | 86.26 |
| Inductive | 2 | 75.15 | 71.27 | 80.85 | 75.76 | 75.15 | 50.91 | 82.59 | 80.63 |
| | 4 | 75.02 | 71.61 | 79.50 | 75.35 | 73.33 | 50.46 | 81.87 | 79.37 |
| | 8 | 73.57 | 71.21 | 75.47 | 73.28 | 70.96 | 47.26 | 80.34 | 78.08 |
| | 16 | 71.57 | 66.98 | 80.44 | 73.10 | 70.92 | 44.31 | 80.40 | 78.49 |
| | 32 | 71.02 | 67.39 | 76.81 | 71.80 | 70.35 | 42.65 | 77.96 | 75.55 |
| | 64 | 68.64 | 63.18 | 83.13 | 71.79 | 70.16 | 39.75 | 77.24 | 74.96 |

**Supplementary Table 7. Hyperparameter tuning results for the SVM on the respective supervision Validation set**. The sizes of the Transductive and Inductive dataset partitions used were 4,619 and 3,099 bioactivity pairs, respectively. All values are reported as percentages. RBF is radial basis function and Polynomial is a third-degree polynomial function.

| Learning | Kernel | Accuracy | Precision | Recall | Micro F1-Score | Macro F1-Score | MCC | AUROC | Average Precision |
|---|---|---|---|---|---|---|---|---|---|
| Transductive | RBF | 70.49 | 82.00 | 70.03 | 75.54 | 76.32 | 39.71 | 77.16 | 84.89 |
| | Linear | 73.28 | 83.71 | 73.19 | 78.10 | 77.75 | 44.91 | 80.20 | 86.88 |
| | Polynomial | 70.86 | 82.05 | 70.69 | 75.95 | 76.51 | 40.21 | 77.85 | 85.25 |
| | Sigmoid | 64.04 | 76.98 | 63.84 | 69.79 | 72.46 | 27.02 | 69.66 | 80.70 |
| Inductive | RBF | 70.96 | 70.68 | 67.54 | 69.07 | 67.96 | 41.76 | 77.61 | 75.65 |
| | Linear | 70.86 | 69.16 | 70.97 | 70.05 | 67.36 | 41.70 | 78.06 | 74.96 |
| | Polynomial | 71.25 | 70.83 | 68.21 | 69.50 | 67.93 | 42.35 | 78.20 | 76.17 |
| | Sigmoid | 66.12 | 64.74 | 64.65 | 64.69 | 63.24 | 32.13 | 72.38 | 71.35 |

**Supplementary Table 8. Hyperparameter tuning results for the XGBoost on the respective supervision Validation set.** The sizes of the Transductive and Inductive dataset partitions used were 4,619 and 3,099 bioactivity pairs, respectively. All values are reported as percentages. Number of Estimators are the maximum number of boosting rounds and Maximum Depth is the maximum depth a tree can have during boosting.

| Learning | Number of Estimators | Maximum Depth | Accuracy | Precision | Recall | Micro F1-Score | Macro F1-Score | MCC | AUROC | Average Precision |
|---|---|---|---|---|---|---|---|---|---|---|
| Transductive | 200 | 2 | 67.24 | 80.53 | 65.50 | 72.24 | 72.27 | 34.38 | 74.46 | 83.46 |
| | 200 | 4 | 71.68 | 83.48 | 70.43 | 76.40 | 76.23 | 42.59 | 79.35 | 86.77 |
| | 200 | 6 | 75.67 | 85.59 | 75.28 | 80.11 | 80.58 | 49.78 | 83.29 | 89.52 |
| | 200 | 8 | 77.77 | 86.34 | 78.21 | 82.07 | 82.12 | 53.44 | 85.92 | 91.32 |
| | 200 | 10 | 78.59 | 87.06 | 78.81 | 82.73 | 82.43 | 55.22 | 87.03 | 92.08 |
| | 200 | 12 | 79.17 | 87.19 | 79.71 | 83.28 | 82.56 | 56.21 | 87.30 | 92.21 |
| | 200 | 14 | 78.98 | 87.67 | 78.78 | 82.99 | 82.41 | 56.24 | 87.30 | 92.24 |
| | 200 | 16 | 78.89 | 87.29 | 79.08 | 82.98 | 82.58 | 55.84 | 87.56 | 92.38 |
| | 200 | 18 | 79.41 | 87.87 | 79.31 | 83.37 | 83.15 | 57.05 | 87.42 | 92.33 |
| | 200 | 20 | 78.96 | 87.67 | 78.74 | 82.97 | 82.82 | 56.20 | 87.26 | 92.29 |
| | 400 | 2 | 69.65 | 82.08 | 68.26 | 74.54 | 75.8 | 38.74 | 76.75 | 84.96 |
| | 400 | 4 | 73.65 | 84.39 | 73.02 | 78.29 | 78.63 | 45.99 | 81.42 | 88.24 |
| | 400 | 6 | 77.01 | 86.27 | 76.91 | 81.32 | 81.27 | 52.26 | 85.05 | 90.73 |

| Learning | Number of Estimators | Maximum Depth | Accuracy | Precision | Recall | Micro F1-Score | Macro F1-Score | MCC | AUROC | Average Precision |
|---|---|---|---|---|---|---|---|---|---|---|
| | 400 | 8 | 79.06 | 87.22 | 79.47 | 83.17 | 83.15 | 56.06 | 87.16 | 92.12 |
| | 400 | 10 | 80.04 | 88.14 | 80.11 | 83.93 | 83.74 | 58.21 | 88.13 | 92.76 |
| | 400 | 12 | 80.08 | 88.01 | 80.34 | 84.00 | 83.73 | 58.19 | 88.32 | 92.89 |
| | 400 | 14 | 79.84 | 88.02 | 79.91 | 83.77 | 83.64 | 57.82 | 88.35 | 92.94 |
| | 400 | 16 | 79.58 | 87.80 | 79.71 | 83.56 | 82.97 | 57.27 | 88.42 | 92.98 |
| | 400 | 18 | 79.95 | 88.15 | 79.94 | 83.85 | 83.53 | 58.08 | 88.32 | 92.85 |
| | 400 | 20 | 79.93 | 87.92 | 80.17 | 83.87 | 83.72 | 57.89 | 88.28 | 92.90 |
| | 600 | 2 | 70.73 | 82.46 | 69.89 | 75.66 | 77.01 | 40.43 | 77.76 | 85.60 |
| | 600 | 4 | 74.60 | 85.13 | 73.89 | 79.11 | 79.37 | 47.92 | 82.43 | 88.92 |
| | 600 | 6 | 78.22 | 87.15 | 78.04 | 82.34 | 82.32 | 54.71 | 85.97 | 91.35 |
| | 600 | 8 | 79.87 | 87.97 | 80.01 | 83.80 | 83.72 | 57.82 | 87.77 | 92.58 |
| | 600 | 10 | 80.43 | 88.39 | 80.51 | 84.26 | 84.26 | 58.98 | 88.57 | 93.07 |
| | 600 | 12 | 80.60 | 88.53 | 80.64 | 84.4 | 84.51 | 59.35 | 88.74 | 93.19 |
| | 600 | 14 | 80.32 | 88.68 | 79.97 | 84.10 | 84.31 | 59.00 | 88.84 | 93.24 |
| | 600 | 16 | 80.15 | 88.41 | 79.97 | 83.98 | 83.80 | 58.56 | 88.81 | 93.22 |
| | 600 | 18 | 80.06 | 88.26 | 80.01 | 83.93 | 83.67 | 58.32 | 88.70 | 93.05 |
| | 600 | 20 | 80.21 | 88.37 | 80.14 | 84.05 | 83.87 | 58.63 | 88.74 | 93.17 |
| | 800 | 2 | 71.21 | 82.71 | 70.49 | 76.11 | 77.04 | 41.27 | 78.44 | 86.04 |
| | 800 | 4 | 75.23 | 85.40 | 74.72 | 79.70 | 79.75 | 49.01 | 83.17 | 89.44 |
| | 800 | 6 | 78.89 | 87.57 | 78.74 | 82.92 | 82.43 | 56.03 | 86.56 | 91.75 |
| | 800 | 8 | 80.19 | 88.28 | 80.21 | 84.05 | 84.05 | 58.54 | 88.16 | 92.87 |
| | 800 | 10 | 80.67 | 88.54 | 80.74 | 84.46 | 84.25 | 59.47 | 88.84 | 93.29 |
| | 800 | 12 | 80.95 | 88.82 | 80.90 | 84.68 | 84.42 | 60.09 | 88.95 | 93.33 |
| | 800 | 14 | 80.73 | 88.98 | 80.34 | 84.44 | 84.62 | 59.85 | 89.05 | 93.4 |
| | 800 | 16 | 80.43 | 88.61 | 80.24 | 84.22 | 84.20 | 59.13 | 89.02 | 93.35 |
| | 800 | 18 | 80.19 | 88.40 | 80.07 | 84.03 | 84.06 | 58.61 | 88.93 | 93.24 |
| | 800 | 20 | 80.34 | 88.54 | 80.17 | 84.15 | 84.03 | 58.95 | 88.99 | 93.35 |
| | 1000 | 2 | 71.34 | 82.90 | 70.49 | 76.20 | 77.27 | 41.62 | 78.98 | 86.4 |
| | 1000 | 4 | 75.77 | 85.62 | 75.45 | 80.21 | 80.24 | 49.96 | 83.75 | 89.87 |
| | 1000 | 6 | 79.24 | 87.75 | 79.14 | 83.23 | 83.04 | 56.70 | 87.02 | 92.04 |
| | 1000 | 8 | 80.41 | 88.38 | 80.47 | 84.24 | 83.83 | 58.95 | 88.38 | 93.01 |
| | 1000 | 10 | 80.82 | 88.80 | 80.71 | 84.56 | 84.44 | 59.87 | 89.03 | 93.45 |
| | 1000 | 12 | 80.97 | 88.80 | 80.97 | 84.71 | 84.41 | 60.11 | 89.12 | 93.49 |
| | 1000 | 14 | 80.99 | 89.18 | 80.57 | 84.66 | 84.96 | 60.39 | 89.19 | 93.50 |
| | 1000 | 16 | 80.60 | 88.67 | 80.47 | 84.37 | 84.35 | 59.44 | 89.15 | 93.46 |
| | 1000 | 18 | 80.47 | 88.48 | 80.47 | 84.29 | 84.40 | 59.11 | 89.09 | 93.33 |
| | 1000 | 20 | 80.45 | 88.59 | 80.31 | 84.24 | 84.30 | 59.15 | 89.13 | 93.45 |

| Learning | Number of Estimators | Maximum Depth | Accuracy | Precision | Recall | Micro F1-Score | Macro F1-Score | MCC | AUROC | Average Precision |
|---|---|---|---|---|---|---|---|---|---|---|
| | 1200 | 2 | 71.99 | 83.36 | 71.16 | 76.78 | 77.49 | 42.87 | 79.41 | 86.70 |
| | 1200 | 4 | 76.01 | 85.76 | 75.72 | 80.42 | 80.50 | 50.41 | 84.26 | 90.22 |
| | 1200 | 6 | 79.63 | 88.23 | 79.27 | 83.51 | 83.10 | 57.62 | 87.42 | 92.29 |
| | 1200 | 8 | 80.69 | 88.52 | 80.81 | 84.49 | 84.21 | 59.49 | 88.60 | 93.17 |
| | 1200 | 10 | 80.82 | 88.91 | 80.57 | 84.54 | 84.25 | 59.94 | 89.15 | 93.54 |
| | 1200 | 12 | 81.12 | 88.97 | 81.04 | 84.82 | 84.54 | 60.46 | 89.22 | 93.57 |
| | 1200 | 14 | 81.08 | 89.22 | 80.67 | 84.73 | 84.99 | 60.55 | 89.30 | 93.57 |
| | 1200 | 16 | 80.82 | 88.83 | 80.67 | 84.55 | 84.52 | 59.89 | 89.25 | 93.52 |
| | 1200 | 18 | 80.58 | 88.53 | 80.61 | 84.38 | 84.27 | 59.32 | 89.20 | 93.41 |
| | 1200 | 20 | 80.67 | 88.77 | 80.47 | 84.42 | 84.33 | 59.61 | 89.21 | 93.50 |
| | 1400 | 2 | 72.44 | 83.65 | 71.66 | 77.19 | 77.88 | 43.73 | 79.76 | 86.96 |
| | 1400 | 4 | 76.51 | 86.12 | 76.18 | 80.85 | 80.85 | 51.41 | 84.72 | 90.53 |
| | 1400 | 6 | 79.76 | 88.22 | 79.51 | 83.64 | 83.32 | 57.82 | 87.78 | 92.51 |
| | 1400 | 8 | 80.82 | 88.60 | 80.94 | 84.60 | 84.24 | 59.75 | 88.73 | 93.26 |
| | 1400 | 10 | 81.10 | 89.11 | 80.84 | 84.77 | 84.26 | 60.52 | 89.24 | 93.60 |
| | 1400 | 12 | 81.16 | 89.06 | 81.00 | 84.84 | 84.67 | 60.59 | 89.31 | 93.64 |
| | 1400 | 14 | 81.03 | 89.21 | 80.61 | 84.69 | 84.77 | 60.48 | 89.36 | 93.62 |
| | 1400 | 16 | 80.90 | 88.96 | 80.67 | 84.61 | 84.58 | 60.11 | 89.31 | 93.56 |
| | 1400 | 18 | 80.60 | 88.62 | 80.54 | 84.38 | 84.19 | 59.41 | 89.28 | 93.48 |
| | 1400 | 20 | 80.62 | 88.82 | 80.34 | 84.37 | 84.37 | 59.57 | 89.27 | 93.54 |
| | 1600 | 2 | 72.70 | 83.70 | 72.09 | 77.46 | 78.04 | 44.11 | 80.07 | 87.19 |
| | 1600 | 4 | 76.66 | 86.10 | 76.48 | 81.01 | 80.81 | 51.62 | 85.13 | 90.79 |
| | 1600 | 6 | 80.02 | 88.36 | 79.81 | 83.87 | 83.50 | 58.32 | 87.98 | 92.64 |
| | 1600 | 8 | 80.97 | 88.80 | 80.97 | 84.71 | 84.18 | 60.11 | 88.86 | 93.34 |
| | 1600 | 10 | 81.25 | 89.22 | 80.97 | 84.90 | 84.53 | 60.83 | 89.32 | 93.66 |
| | 1600 | 12 | 81.21 | 89.07 | 81.07 | 84.88 | 84.62 | 60.67 | 89.38 | 93.69 |
| | 1600 | 14 | 81.19 | 89.24 | 80.84 | 84.83 | 84.87 | 60.74 | 89.41 | 93.65 |
| | 1600 | 16 | 80.95 | 88.97 | 80.74 | 84.65 | 84.59 | 60.18 | 89.37 | 93.61 |
| | 1600 | 18 | 80.62 | 88.62 | 80.57 | 84.40 | 84.35 | 59.45 | 89.34 | 93.51 |
| | 1600 | 20 | 80.78 | 88.91 | 80.51 | 84.50 | 84.46 | 59.87 | 89.33 | 93.58 |
| | 1800 | 2 | 73.07 | 84.07 | 72.32 | 77.75 | 78.40 | 44.92 | 80.32 | 87.38 |
| | 1800 | 4 | 77.12 | 86.54 | 76.78 | 81.37 | 81.07 | 52.62 | 85.50 | 91.03 |
| | 1800 | 6 | 80.02 | 88.45 | 79.71 | 83.85 | 83.34 | 58.37 | 88.15 | 92.75 |
| | 1800 | 8 | 81.03 | 88.90 | 80.97 | 84.75 | 84.23 | 60.28 | 88.95 | 93.39 |
| | 1800 | 10 | 81.27 | 89.20 | 81.04 | 84.92 | 84.69 | 60.85 | 89.39 | 93.71 |
| | 1800 | 12 | 81.32 | 89.23 | 81.07 | 84.96 | 84.73 | 60.94 | 89.45 | 93.74 |
| | 1800 | 14 | 81.36 | 89.27 | 81.10 | 84.99 | 84.99 | 61.03 | 89.45 | 93.68 |

| Learning | Number of Estimators | Maximum Depth | Accuracy | Precision | Recall | Micro F1-Score | Macro F1-Score | MCC | AUROC | Average Precision |
|---|---|---|---|---|---|---|---|---|---|---|
| | 1800 | 16 | 80.86 | 88.86 | 80.71 | 84.59 | 84.40 | 59.98 | 89.42 | 93.63 |
| | 1800 | 18 | 80.69 | 88.69 | 80.61 | 84.45 | 84.45 | 59.59 | 89.38 | 93.54 |
| | 1800 | 20 | 80.82 | 88.91 | 80.57 | 84.54 | 84.57 | 59.94 | 89.39 | 93.63 |
| | 2000 | 2 | 73.13 | 83.90 | 72.65 | 77.87 | 78.56 | 44.87 | 80.57 | 87.57 |
| | 2000 | 4 | 77.33 | 86.67 | 77.01 | 81.56 | 81.29 | 53.04 | 85.77 | 91.20 |
| | 2000 | 6 | 80.17 | 88.50 | 79.91 | 83.99 | 83.44 | 58.65 | 88.32 | 92.86 |
| | 2000 | 8 | 81.08 | 89.02 | 80.90 | 84.77 | 84.16 | 60.42 | 89.05 | 93.46 |
| | 2000 | 10 | 81.27 | 89.26 | 80.97 | 84.91 | 84.56 | 60.89 | 89.41 | 93.73 |
| | 2000 | 12 | 81.42 | 89.31 | 81.17 | 85.05 | 84.69 | 61.16 | 89.50 | 93.76 |
| | 2000 | 14 | 81.40 | 89.34 | 81.10 | 85.02 | 85.01 | 61.14 | 89.49 | 93.72 |
| | 2000 | 16 | 81.01 | 89.01 | 80.81 | 84.71 | 84.40 | 60.31 | 89.44 | 93.65 |
| | 2000 | 18 | 80.73 | 88.67 | 80.71 | 84.50 | 84.49 | 59.65 | 89.42 | 93.57 |
| | 2000 | 20 | 80.97 | 89.06 | 80.67 | 84.66 | 84.51 | 60.28 | 89.42 | 93.65 |
| | 200 | 2 | 67.51 | 67.44 | 62.50 | 64.88 | 62.96 | 34.81 | 74.06 | 70.68 |
| | 200 | 4 | 71.25 | 70.83 | 68.21 | 69.50 | 68.35 | 42.35 | 77.88 | 75.47 |
| | 200 | 6 | 73.25 | 72.61 | 71.10 | 71.85 | 70.31 | 46.38 | 80.42 | 78.68 |
| | 200 | 8 | 73.73 | 72.59 | 72.78 | 72.68 | 70.09 | 47.39 | 81.78 | 80.77 |
| | 200 | 10 | 75.22 | 73.90 | 74.80 | 74.35 | 71.98 | 50.38 | 82.65 | 81.98 |
| | 200 | 12 | 74.96 | 73.54 | 74.73 | 74.13 | 71.57 | 49.88 | 82.30 | 81.54 |
| | 200 | 14 | 74.28 | 72.51 | 74.80 | 73.64 | 71.20 | 48.57 | 82.04 | 81.69 |
| | 200 | 16 | 74.09 | 72.11 | 75.07 | 73.56 | 71.21 | 48.21 | 81.78 | 81.52 |
| | 200 | 18 | 73.06 | 71.00 | 74.19 | 72.56 | 69.89 | 46.16 | 81.28 | 81.05 |
| Inductive | 200 | 20 | 73.67 | 71.88 | 74.19 | 73.02 | 69.84 | 47.34 | 80.95 | 80.51 |
| | 400 | 2 | 69.73 | 69.50 | 65.86 | 67.63 | 65.63 | 39.30 | 76.16 | 73.69 |
| | 400 | 4 | 72.57 | 71.70 | 70.83 | 71.26 | 69.90 | 45.03 | 79.45 | 77.41 |
| | 400 | 6 | 74.22 | 73.29 | 72.85 | 73.07 | 71.59 | 48.34 | 81.85 | 80.52 |
| | 400 | 8 | 75.19 | 74.05 | 74.40 | 74.22 | 71.29 | 50.30 | 83.24 | 82.45 |
| | 400 | 10 | 76.02 | 74.69 | 75.74 | 75.21 | 73.26 | 52.00 | 84.01 | 83.50 |
| | 400 | 12 | 75.73 | 74.47 | 75.27 | 74.87 | 72.88 | 51.41 | 83.73 | 83.16 |
| | 400 | 14 | 76.44 | 74.67 | 77.08 | 75.86 | 73.22 | 52.90 | 83.68 | 83.46 |
| | 400 | 16 | 75.99 | 74.31 | 76.41 | 75.35 | 73.18 | 51.98 | 83.63 | 83.38 |
| | 400 | 18 | 75.09 | 73.55 | 75.13 | 74.34 | 71.35 | 50.15 | 83.38 | 83.18 |
| | 400 | 20 | 74.51 | 72.81 | 74.87 | 73.82 | 71.01 | 49.01 | 83.13 | 82.85 |
| | 600 | 2 | 69.96 | 69.43 | 66.87 | 68.13 | 66.78 | 39.76 | 77.02 | 74.87 |
| | 600 | 4 | 73.54 | 72.57 | 72.18 | 72.37 | 71.05 | 46.99 | 80.37 | 78.62 |
| | 600 | 6 | 74.60 | 73.54 | 73.59 | 73.56 | 71.61 | 49.13 | 82.40 | 81.35 |
| | 600 | 8 | 76.28 | 75.18 | 75.54 | 75.36 | 72.69 | 52.50 | 83.76 | 83.13 |

| Learning | Number of Estimators | Maximum Depth | Accuracy | Precision | Recall | Micro F1-Score | Macro F1-Score | MCC | AUROC | Average Precision |
|---|---|---|---|---|---|---|---|---|---|---|
| | 600 | 10 | 76.83 | 75.74 | 76.14 | 75.94 | 73.87 | 53.60 | 84.61 | 84.18 |
| | 600 | 12 | 76.31 | 75.03 | 75.94 | 75.48 | 73.46 | 52.58 | 84.34 | 83.92 |
| | 600 | 14 | 76.51 | 74.97 | 76.68 | 75.81 | 73.30 | 53.00 | 84.27 | 84.06 |
| | 600 | 16 | 76.83 | 75.50 | 76.61 | 76.05 | 74.11 | 53.62 | 84.21 | 84.00 |
| | 600 | 18 | 75.86 | 74.57 | 75.47 | 75.02 | 72.30 | 51.68 | 83.98 | 83.89 |
| | 600 | 20 | 75.22 | 73.81 | 75.00 | 74.40 | 72.26 | 50.39 | 83.83 | 83.64 |
| | 800 | 2 | 70.28 | 69.62 | 67.61 | 68.60 | 66.15 | 40.42 | 77.50 | 75.69 |
| | 800 | 4 | 73.54 | 72.69 | 71.91 | 72.30 | 70.81 | 46.98 | 80.83 | 79.18 |
| | 800 | 6 | 74.38 | 73.04 | 73.92 | 73.48 | 71.73 | 48.70 | 82.77 | 81.87 |
| | 800 | 8 | 76.35 | 75.32 | 75.47 | 75.39 | 71.96 | 52.62 | 84.13 | 83.53 |
| | 800 | 10 | 77.22 | 76.31 | 76.21 | 76.26 | 74.04 | 54.36 | 84.88 | 84.47 |
| | 800 | 12 | 76.90 | 75.63 | 76.55 | 76.09 | 73.94 | 53.74 | 84.68 | 84.27 |
| | 800 | 14 | 76.61 | 75.32 | 76.28 | 75.79 | 72.96 | 53.16 | 84.52 | 84.33 |
| | 800 | 16 | 76.70 | 75.47 | 76.28 | 75.87 | 74.19 | 53.35 | 84.48 | 84.30 |
| | 800 | 18 | 75.90 | 74.82 | 75.07 | 74.94 | 72.66 | 51.72 | 84.27 | 84.20 |
| | 800 | 20 | 75.64 | 74.26 | 75.40 | 74.82 | 72.52 | 51.23 | 84.19 | 83.99 |
| | 1000 | 2 | 71.09 | 70.50 | 68.41 | 69.44 | 66.87 | 42.04 | 77.93 | 76.25 |
| | 1000 | 4 | 73.60 | 72.60 | 72.31 | 72.46 | 70.93 | 47.12 | 81.15 | 79.64 |
| | 1000 | 6 | 74.86 | 73.59 | 74.33 | 73.96 | 72.31 | 49.67 | 82.99 | 82.13 |
| | 1000 | 8 | 76.51 | 75.50 | 75.60 | 75.55 | 72.32 | 52.95 | 84.34 | 83.79 |
| | 1000 | 10 | 77.15 | 76.14 | 76.34 | 76.24 | 74.45 | 54.24 | 85.07 | 84.68 |
| | 1000 | 12 | 76.93 | 75.82 | 76.28 | 76.05 | 73.75 | 53.79 | 84.86 | 84.49 |
| | 1000 | 14 | 76.67 | 75.31 | 76.48 | 75.89 | 73.26 | 53.30 | 84.70 | 84.51 |
| | 1000 | 16 | 76.70 | 75.43 | 76.34 | 75.89 | 74.27 | 53.36 | 84.64 | 84.50 |
| | 1000 | 18 | 76.12 | 75.03 | 75.34 | 75.18 | 72.84 | 52.18 | 84.46 | 84.40 |
| | 1000 | 20 | 75.80 | 74.47 | 75.47 | 74.97 | 72.79 | 51.55 | 84.38 | 84.20 |
| | 1200 | 2 | 71.60 | 70.88 | 69.35 | 70.11 | 68.56 | 43.08 | 78.22 | 76.56 |
| | 1200 | 4 | 73.86 | 72.75 | 72.85 | 72.80 | 71.18 | 47.65 | 81.39 | 80.02 |
| | 1200 | 6 | 75.19 | 73.92 | 74.66 | 74.29 | 72.34 | 50.31 | 83.14 | 82.32 |
| | 1200 | 8 | 76.83 | 75.70 | 76.21 | 75.95 | 72.98 | 53.60 | 84.55 | 84.02 |
| | 1200 | 10 | 77.15 | 76.21 | 76.21 | 76.21 | 74.08 | 54.24 | 85.20 | 84.88 |
| | 1200 | 12 | 77.02 | 75.90 | 76.41 | 76.16 | 74.15 | 53.99 | 85.01 | 84.68 |
| | 1200 | 14 | 76.96 | 75.53 | 76.95 | 76.23 | 73.60 | 53.89 | 84.79 | 84.61 |
| | 1200 | 16 | 77.12 | 75.91 | 76.68 | 76.30 | 74.68 | 54.19 | 84.74 | 84.61 |
| | 1200 | 18 | 76.51 | 75.47 | 75.67 | 75.57 | 73.50 | 52.95 | 84.60 | 84.52 |
| | 1200 | 20 | 76.06 | 74.73 | 75.74 | 75.23 | 72.95 | 52.07 | 84.53 | 84.35 |
| | 1400 | 2 | 72.02 | 71.19 | 70.09 | 70.64 | 69.10 | 43.93 | 78.51 | 76.76 |

| Learning | Number of Estimators | Maximum Depth | Accuracy | Precision | Recall | Micro F1-Score | Macro F1-Score | MCC | AUROC | Average Precision |
|---|---|---|---|---|---|---|---|---|---|---|
| | 1400 | 4 | 73.89 | 72.77 | 72.92 | 72.84 | 70.95 | 47.71 | 81.57 | 80.30 |
| | 1400 | 6 | 75.35 | 74.04 | 74.93 | 74.48 | 72.97 | 50.64 | 83.34 | 82.57 |
| | 1400 | 8 | 76.64 | 75.40 | 76.21 | 75.80 | 73.06 | 53.22 | 84.69 | 84.18 |
| | 1400 | 10 | 77.06 | 76.02 | 76.28 | 76.15 | 74.29 | 54.05 | 85.28 | 85.02 |
| | 1400 | 12 | 77.38 | 76.22 | 76.88 | 76.55 | 75.27 | 54.71 | 85.10 | 84.78 |
| | 1400 | 14 | 77.15 | 75.83 | 76.95 | 76.38 | 73.91 | 54.27 | 84.87 | 84.69 |
| | 1400 | 16 | 77.25 | 76.08 | 76.75 | 76.41 | 74.36 | 54.45 | 84.84 | 84.71 |
| | 1400 | 18 | 76.57 | 75.47 | 75.87 | 75.67 | 73.65 | 53.08 | 84.71 | 84.62 |
| | 1400 | 20 | 76.51 | 75.40 | 75.81 | 75.60 | 73.27 | 52.95 | 84.65 | 84.46 |
| | 1600 | 2 | 71.96 | 71.16 | 69.96 | 70.55 | 69.16 | 43.80 | 78.78 | 77.02 |
| | 1600 | 4 | 73.83 | 72.67 | 72.92 | 72.79 | 71.23 | 47.58 | 81.77 | 80.54 |
| | 1600 | 6 | 75.54 | 74.24 | 75.13 | 74.68 | 72.90 | 51.03 | 83.45 | 82.79 |
| | 1600 | 8 | 76.73 | 75.52 | 76.28 | 75.89 | 72.86 | 53.42 | 84.77 | 84.31 |
| | 1600 | 10 | 77.28 | 76.20 | 76.61 | 76.41 | 74.67 | 54.50 | 85.34 | 85.13 |
| | 1600 | 12 | 77.54 | 76.33 | 77.15 | 76.74 | 75.41 | 55.03 | 85.17 | 84.83 |
| | 1600 | 14 | 77.19 | 75.91 | 76.88 | 76.39 | 74.14 | 54.33 | 84.95 | 84.77 |
| | 1600 | 16 | 77.06 | 75.85 | 76.61 | 76.23 | 74.40 | 54.06 | 84.94 | 84.83 |
| | 1600 | 18 | 76.57 | 75.47 | 75.87 | 75.67 | 73.51 | 53.08 | 84.78 | 84.67 |
| | 1600 | 20 | 76.51 | 75.44 | 75.74 | 75.59 | 73.06 | 52.95 | 84.71 | 84.52 |
| | 1800 | 2 | 72.35 | 71.74 | 69.96 | 70.84 | 69.39 | 44.56 | 78.98 | 77.22 |
| | 1800 | 4 | 73.96 | 72.68 | 73.32 | 73.00 | 71.38 | 47.86 | 81.95 | 80.73 |
| | 1800 | 6 | 75.73 | 74.44 | 75.34 | 74.88 | 73.03 | 51.42 | 83.60 | 82.92 |
| | 1800 | 8 | 76.77 | 75.60 | 76.21 | 75.90 | 73.32 | 53.48 | 84.83 | 84.45 |
| | 1800 | 10 | 77.57 | 76.56 | 76.81 | 76.69 | 75.03 | 55.08 | 85.40 | 85.21 |
| | 1800 | 12 | 77.35 | 76.10 | 77.02 | 76.55 | 75.11 | 54.65 | 85.20 | 84.88 |
| | 1800 | 14 | 77.22 | 76.03 | 76.75 | 76.39 | 74.13 | 54.38 | 85.02 | 84.83 |
| | 1800 | 16 | 77.06 | 75.88 | 76.55 | 76.21 | 74.25 | 54.06 | 85.00 | 84.86 |
| | 1800 | 18 | 76.83 | 75.74 | 76.14 | 75.94 | 73.78 | 53.60 | 84.82 | 84.71 |
| | 1800 | 20 | 76.41 | 75.42 | 75.47 | 75.45 | 73.21 | 52.75 | 84.76 | 84.55 |
| | 2000 | 2 | 72.35 | 71.62 | 70.23 | 70.92 | 69.43 | 44.57 | 79.13 | 77.38 |
| | 2000 | 4 | 74.02 | 72.84 | 73.19 | 73.01 | 71.18 | 47.98 | 82.07 | 80.91 |
| | 2000 | 6 | 75.99 | 74.54 | 75.94 | 75.23 | 73.09 | 51.95 | 83.77 | 83.14 |
| | 2000 | 8 | 76.93 | 75.55 | 76.81 | 76.17 | 73.32 | 53.82 | 84.92 | 84.53 |
| | 2000 | 10 | 77.70 | 76.69 | 76.95 | 76.82 | 74.91 | 55.34 | 85.43 | 85.22 |
| | 2000 | 12 | 77.44 | 76.25 | 77.02 | 76.63 | 75.21 | 54.84 | 85.27 | 84.95 |
| | 2000 | 14 | 77.28 | 76.03 | 76.95 | 76.49 | 74.59 | 54.52 | 85.07 | 84.88 |
| | 2000 | 16 | 77.15 | 75.93 | 76.75 | 76.34 | 74.64 | 54.26 | 85.06 | 84.90 |

| **Learning** | **Number of Estimators** | **Maximum Depth** | **Accuracy** | **Precision** | **Recall** | **Micro F1-Score** | **Macro F1-Score** | **MCC** | **AUROC** | **Average Precision** |
|---|---|---|---|---|---|---|---|---|---|---|
| | 2000 | 18 | 76.83 | 75.67 | 76.28 | 75.97 | 73.72 | 53.61 | 84.86 | 84.76 |
| | 2000 | 20 | 76.44 | 75.40 | 75.60 | 75.50 | 73.42 | 52.82 | 84.79 | 84.59 |
